# ORP1L regulates dynein clustering on endolysosmal membranes in response to cholesterol levels

**DOI:** 10.1101/2020.08.28.273037

**Authors:** Shreyasi Thakur, Peter K. Relich, Elena M. Sorokina, Melina T. Gyparaki, Melike Lakadamyali

## Abstract

The sub-cellular positioning of endolysosomes is crucial for regulating their function. Particularly, the positioning of endolysosomes between the cell periphery versus the peri-nuclear region impacts autophagy, mTOR (mechanistic target of rapamycin) signaling and other processes. The mechanisms that regulate the positioning of endolysosomes at these two locations are still being uncovered. Here, using super-resolution microscopy, we show that the retrograde motor dynein forms nano-clusters on endolysosomal membranes containing 1-2 dyneins, with an average of ~3 nanoclusters per endolysosome. These data suggest that a very small number of dynein motors (1-6) drive endolysosome motility. Surprisingly, dynein nano-clusters are slightly larger on peripheral endolysosomes having higher cholesterol levels compared to peri-nuclear ones. By perturbing endolysosomal membrane cholesterol levels, we show that dynein copy number within nano-clusters is influenced by the amount of endolysosomal cholesterol while the total number of nano-clusters per endolysosome is independent of cholesterol. Finally, we show that the dynein adapter protein ORP1L (Oxysterol Binding Protein Homologue) regulates the number of dynein motors within nano-clusters in response to cholesterol levels. We propose a new model by which endolysosomal transport and positioning is influenced by the cholesterol sensing adapter protein ORP1L, which influences dynein’s copy number within nano-clusters.

## Introduction

Late endosomes, lysosomes and autolysosomes constitute a broad class of sub-cellular compartments (Klumperman and Raposo, 2014; Wijdeven et al., 2016) that we will refer to here as endolysosomes for simplicity. Endolysosmal compartments play key cellular roles including transport of cellular proteins destined for degradation, metabolic sensing, membrane repair and signaling (Gould and Lippincott-Schwartz, 2009).The maturation level, fusion capacity with other sub-cellular compartments and downstream function of endolysosomes are regulated by their intracellular transport and sub-cellular positioning. For example, dispersal of lysosomes from the peri-nuclear region to the cell periphery increases their association with mTORC1 (mechanistic target of rapamycin complex 1) and leads to downregulation of autophagy (Kimura et al., 2008; Korolchuk and Rubinsztein, 2011; Korolchuk et al., 2011; Pu et al., 2016). In addition, it is known that endolysosmes have heterogeneity in their membrane lipid composition, which influences their function. In general, different cellular organelles have unique lipid composition and there is a lipid gradient from ER to the plasma membrane with highest amount of cholesterol and sphingolipids present in the plasma membrane (Abdul-Hammed et al., 2010; Brugger et al., 2000). Spatially compartmentalized endolysosomal populations also have distinct membrane lipid composition, resulting in varying membrane proteomics and lipid-protein interplay, effecting their maturation and functions. For example, early endosomes mostly located peripherally have higher membrane phosphatidylinositol 3-phosphate (PI(3)P) and cholesterol content (Arumugam and Kaur, 2017; Bissig and Gruenberg, 2013; Huotari and Helenius, 2011) and are rich in PI(3)P-binding proteins which regulates lysosome targeting and retrograde transport (Bissig and Gruenberg, 2013; Lemmon, 2008). More peri-nuclearly enriched late endosomes, on the other hand, have limiting membranes with low phosphatidylserine, cholesterol, and sphingomyelin content, resulting in distinct membrane dynamics, proteomics and functions (Kobayashi et al., 2002; Redpath et al., 2020).

The transport and subcellular positioning of endolysosomes are in turn regulated by a myriad of mechanisms (Maday et al., 2014) including motor activation (Elshenawy et al., 2020; Fu and Holzbaur, 2014; Fu et al., 2014), motor tug-of-war (Belyy et al., 2016; Hendricks et al., 2010; Soppina et al., 2009) and association of motors with microtubule tracks having distinct post-translational modifications (Guardia et al., 2016; Mohan et al., 2019; Nirschl et al., 2016). Peripheral transport of endolysosomes is mediated by kinesin motors belonging to different kinesin families (Kif5, Kif1, Kif3) (Brown et al., 2005; Cardoso et al., 2009; Encalada et al., 2011; Mohan et al., 2019; Rosa-Ferreira and Munro, 2011), whereas dynein is responsible for their retrograde transport toward the perinuclear region (Granger et al., 2014; Reck-Peterson et al., 2018). Recent discovery of dynein activating adapters have revolutionized our understanding of how dynein mediates efficient retrograde transport (Elshenawy et al., 2020; McKenney et al., 2014; Olenick and Holzbaur, 2019; Reck-Peterson et al., 2018; Schroeder and Vale, 2016). Dynein assembles into an autoinhibitory, weakly processive conformation, while dynein activating adaptors are crucial for dynein’s assembly with dynactin and the processive motility of the dynein-dynactin complex on microtubules (Chowdhury et al., 2015; McKenney et al., 2014; Schroeder and Vale, 2016; Urnavicius et al., 2018). Recent Cryo-EM studies showed that certain early endosomal dynein activating adapters including BICD2 and Hook3 recruit two dynein dimers (Urnavicius et al., 2018). *In vitro* assays further showed that these heteromeric dynein complexes move faster and navigate obstacles better compared to single dynein (Elshenawy et al., 2019; Ferro et al., 2019; Urnavicius et al., 2018). Hence, in addition to mediating dynein’s interaction with dynactin and bringing dynein out of its auto-inhibitory confirmation, these activators further improve the efficiency of dynein mediated motility by allowing two copies of dynein to assemble together into complexes. Yet, to date, the stoichiometry of adapter-dynein-dynactin complexes on sub-cellular compartments i*n vivo* and how these complexes are spatially organized on the membrane of sub-cellular compartments are not known. Dynein is recruited to endolysosomal compartments via the cholesterol-sensing tripartite complex Rab7-RILP-ORP1L (Johansson et al., 2007; Pfeffer, 2001; Rocha et al., 2009). Whether this tripartite complex can recruit multiple copies of dynein dimers to improve the efficiency of retrograde transport of endolysosomal compartments in response to cellular cues is unknown.

Super-resolution is a powerful tool for studying spatial nano-organization of proteins within the cell, yet, only a handful of studies have been carried out to date to visualize proteins on the membrane of sub-cellular compartments (Franke et al., 2019; Puchner et al., 2013). Previously, using super-resolution microscopy, we showed that dynein forms nano-clusters on microtubules consisting of small teams of dynein motors (Cella Zanacchi et al., 2019; Zanacchi et al., 2017). However, whether these nano-clusters are formed on the membrane of endolysosomes, the mechanisms of nano-cluster formation and whether formation of larger nano-clusters containing more dynein motors lead to more efficient retrograde transport are not known. Here, using quantitative super-resolution microscopy, we show that dynein resides in on average ~3 nano-clusters on endolysosomes, consisting of 1-2 dyneins, suggesting that on average a very small total number of dynein motors (1-6) drive endolysosomal transport. Our data further suggest that an increase in the percentage of dynein multimers present within the nano-clusters on endolysosomes correlates with longer run lengths and faster velocity in the retrograde direction as well as an increased peri-nuclear clustering of endolysosomes. The copy number of dynein within nano-clusters is in turn regulated by membrane cholesterol levels and ORP1-L’s cholesterol sensing domain.

## Results

### Dynein forms nano-clusters on endolysosomes containing 1-2 dynein motors with a switch to larger nano-clusters on peripheral versus peri-nuclear endolysosomes

To visualize the spatial organization of dynein on endolysosomal membranes we expressed ORP1L WT-mCherry in HeLa cells in which endogenous ORP1L was targeted by CRISPR/Cas9 mutagenesis (ORP1L-KO) (Zhao and Ridgway, 2017). The ORP1L-mCherry signal substantially overlapped with that of CD63, an endolysosomal marker (Beatty, 2006; van der Kant et al., 2013; Vanlandingham and Ceresa, 2009), indicating that ORP1L marks endolysosomes (Figure S1A). Hence, we took ORP1L WT-mCherry positive sub-compartments in the wide-field images that were isolated, round and diffraction limited in size (200-500 nm FWHM) as an endolysosome for further analysis (Figure 1A and Figure S1B). To determine whether dynein was clustered on these endolysosomes, we carried out super-resolution imaging of dynein labeled with an antibody against the dynein intermediate chain (IC74). We validated the IC74 antibody using a HeLa cell line stably expressing IC74 tagged with GFP(Kobayashi et al., 2002) (Figure S1C). There was high co-localization between the GFP signal and the IC74 antibody (Manders coefficient 0.8 for IC74 antibody colocalizing with GFP) showing that the IC74 antibody had high labeling specificity and efficiency. Super-resolution images of IC74 revealed nano-clusters within the cell cytoplasm similar to what we have previously demonstrated (Cella Zanacchi et al., 2019) (Figure 1A-B). We manually cropped the central intensity peak of ORP1L WT-mCherry positive endolysosomes (Figure S1B and Methods) and used it as a mask in the super-resolution image to specifically segment dynein nano-clusters that overlapped with an endolysosomal compartment (Figure 1A-B). Dynein super-resolution images were further segmented into individual nano-clusters using a previously developed Voronoi tessellation approach (Figure S1D) (Levet et al., 2015). We then quantified the number of dynein nano-clusters as well as the number of localizations per dynein nano-cluster for peripherally- and peri-nuclearly-positioned endolysosomes (Figure S1E and Figure 1C). The peripheral and peri-nuclear endolysosomes were separated manually based on their proximity to the cell nucleus (Figure S1F). Endolysosomes contained on average ~2 dynein nano-clusters and there was no difference in the number of dynein nano-clusters associated to peripheral or perinuclear endolysosomes (Figure S1E). The number of localizations per nano-cluster is proportional to the nano-cluster size as well as to the copy number of dynein within nano-clusters (Cella Zanacchi et al., 2019; Zanacchi et al., 2017). Surprisingly, we found that the dynein nano-clusters associated to peripheral endolysosomes contained a significantly higher number of localizations compared to peri-nuclear endolysosomes (Figure 1C, mean = 67±1.9 for peripheral and 54±1.8 for perinuclear endolysosomes).

**Figure 1:**
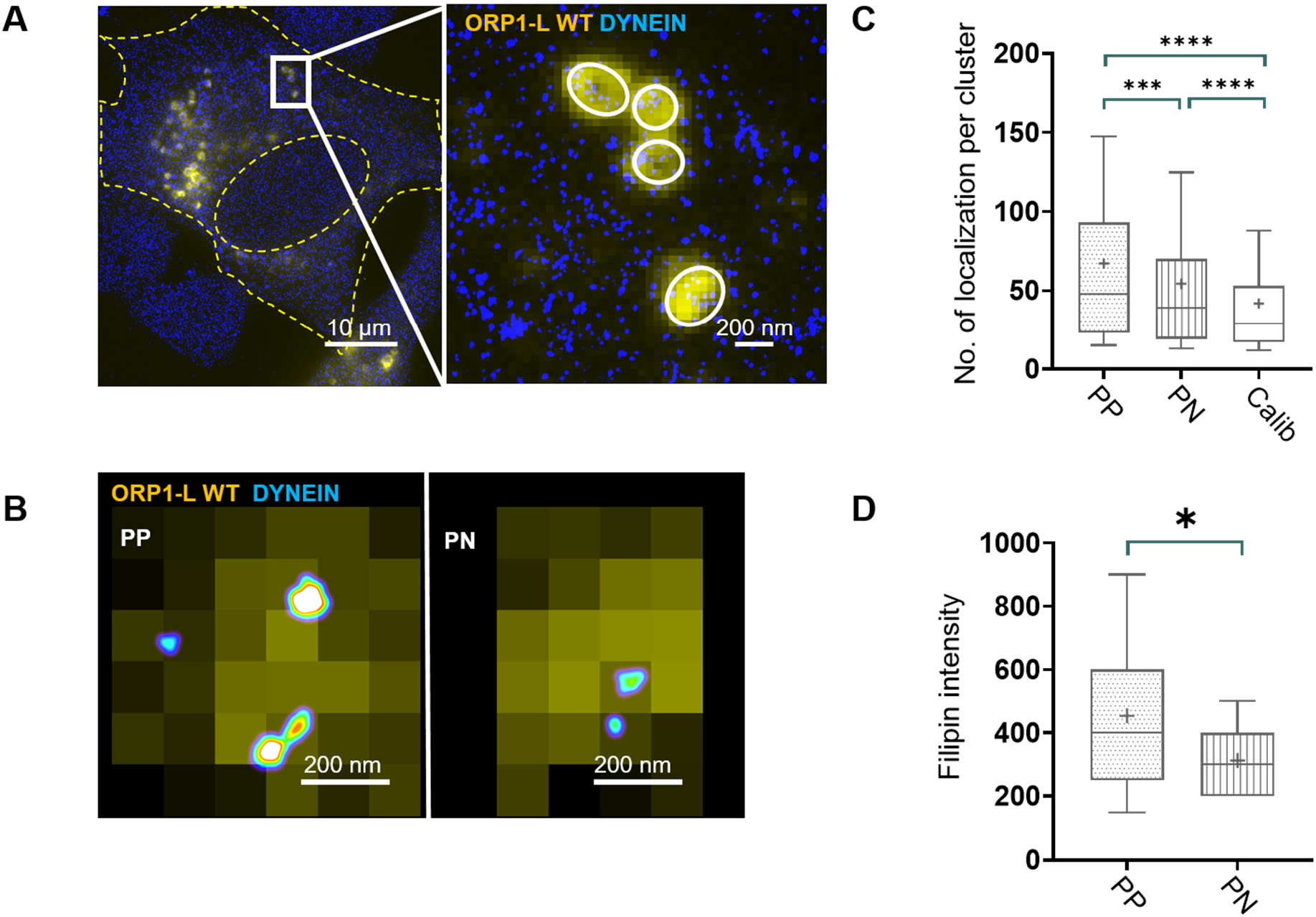
Dynein forms larger nano-clusters containing a higher proportion of dynein multimers on endolysosomes positioned at the cell periphery: (A) Cropped wide-field image of cells expressing full length ORP1L fused to mCherry (ORP1L WT-mCherry, yellow) overlaid with super-resolution image of dynein (blue). Cell edge and nucleus are highlighted in yellow dashed lines. A zoom of the white rectangle is shown in which ORP1L WT-mCherry positive endolysosomes are highlighted with white circles. (B) An overlay of wide-field image of ORP1L WT-mCherry (yellow) and super-resolution image of immunostained dynein (image is color coded according to localization density with higher density corresponding to white and lower density corresponding to cyan) for an endolysosome positioned at the cell periphery (PP) and peri-nuclear region (PN) in Hela ORP1L-KO cells (C) Box plot showing the number of localizations per dynein nano-cluster for peripherally positioned endolysosomes (PP) (n=195 endolysosomes from n=6 cells, n=2 experiments, mean 67±1.9), peri-nuclearly positioned endolysosomes (PN) (n=210 endolysosomes from n=6 cells, n=2 experiments, mean 54±1.8) in ORP1L WT-mCherry expressing Hela ORPiL-KO cells and for dynein that has been labeled with a 100-fold dilution of the primary antibody for sparsely labeling single IC74 subunits in Hela WT cells (Calib) (n=6 cells, n=1 experiments, see methods, mean 42±0.2). The box corresponds to 25-75 percentile, the line corresponds to the median, the cross corresponds to the mean, and the whiskers correspond to 10-90 percentile. Statistical significance was assessed using a Kolmogorov-Smirnov-test with a p-value of. ***: 0.001, ****: <0.0001. (D) Box plot showing the intensity of filipin, which binds cholesterol, on peripherally positioned endolysosomes (PP) (n=44 endolysosomes from n=4 cells, n=2 experiments, mean 454±40) versus peri-nuclearly positioned endolysosomes (PN) (n=41 endolysosomes from n=5 cells, n=2 experiments, mean 312±19) in ORP1L WT-mCherry expressing Hela ORP1L-KO cells. The box corresponds to 25-75 percentile, the line corresponds to the median, the cross corresponds to the mean and the whiskers correspond to 10-90 percentile. Statistical significance was assessed using a Kolmogorov-Smirnov-test with a p-value of 0.02

To obtain a quantitative estimate of the copy number of dynein within nano-clusters, we carried out a calibration experiment in which we diluted the primary antibody by 100-fold, aiming to sparsely label single copies of the dynein IC74 subunit (Ehmann et al., 2014) (Figure S1G and Methods). We then used the number of localizations per nano-cluster in these sparse labeling experiments as a calibration corresponding to single IC74 subunit (Figure 1C, Figure S1H). We previously showed that fitting the distribution of number of localizations per nano-cluster to a linear convolution of monomeric calibration functions (*f_1_, f_2_, f_3_*…) enables estimation of the copy number composition of a protein of interest in super-resolution images (Zanacchi et al., 2017). We used this approach and stopped the fitting when the residuals for the fit stopped changing, which corresponded to *f_3_* for both peripheral and perinuclear endolysosomes (see Figure S1H for residuals when fitting to N=1, N=2, N=3 and N=4). Using this approach, we estimated that nano-clusters on peri-nuclear endolysosomes consist of 76% single IC74, 4% 2 IC74s, 20% 3 IC74s and those on peripheral endolysosomes consist of 69% single IC74, 0.1% 2 IC74s, 31% 3 IC74s (Figure S1H). Given that each dynein motor is a dimer of IC74, the presence of single IC74 subunits in the quantification is likely due to steric effects associated with labeling two subunits of the same protein in close proximity. We thus took the monomeric IC74 weights in the fit to represent a partially labeled single dynein motor. Further, the weights corresponding to trimeric IC74 represent partially labelled dynein multimers, most likely consisting of 2 copies of dynein. It is possible that the dimeric IC74 weight in the fit corresponds partially to a single dynein and partially to dynein multimers. Since we could not clearly assign this population to a given category (single versus multiple dynein) and since this population was a minority (0.1% and 4% on peripheral and perinuclear endolysosomes, respectively) and would not affect our conclusions, we excluded it from consideration. These data suggest that majority of dynein is present in single copy within the nano-clusters and a small proportion of nanoclusters contain dynein multimers, likely consisting of two copies of dynein. Interestingly, compared to peri-nuclear endolysosomes, on peripheral endolysosomes the proportion of nano-clusters containing dynein multimers increased by about 1.6-fold (from 20% to 31%) (Figure S1H). These differences in the dynein clustering were not due to differences in the imaging depth of peripheral versus peri-nuclear endolysosomes since cytosolic clusters not associated to endolysosomes analyzed from the same peripheral and peri-nuclear regions showed no differences in the number of localizations per nano-cluster (Figure S1I). It is important to mention that the proportion of multimeric dynein is likely underestimated in these quantifications due to the steric hindrance problems in labeling multiple copies of IC74 in close proximity and this problem will be exacerbated under conditions in which endolysosomes contain a higher proportion of dynein multimers. Hence, the difference in the proportion of dynein multimers between peripheral and perinuclear endolysosomes is likely larger than our quantitative estimate. Overall, given that each endolysosome contains on average ~3 dynein nano-clusters independent of its position within the cell, and the nano-clusters are mainly composed of 1-2 dynein motors, these data suggest that a small total number of dynein motors (1-6) drive the motility of endolysosomes. These results are consistent with previous estimates using single step photobleaching and western blot analysis on lysosomal compartments (Hendricks et al., 2010).

It is known that positioning of endolysosomal compartments in the cell affects their membrane composition, maturation and signaling (Cabukusta and Neefjes, 2018; Hu et al., 2015; Hyttinen et al., 2013). We thus asked if the membrane cholesterol content may be responsible for the increase in the presence of 2 copies of dynein within the nano-clusters on peripheral endolysosomes. To start addressing this question, we measured the cholesterol levels of peripheral and peri-nuclear endolysosomes by labeling ORP1L WT-mCherry expressing HeLa cells with filipin, a toxin that binds cholesterol. We measured the filipin intensity on ORP1L WT-mCherry positive endolysosomes and found that membrane cholesterol levels of peripheral endolysosomes (mean intensity = 454±40 a.u.) were indeed higher compared to peri-nculear endolysosomes (mean intensity = 312±19 a.u.) (Figure 1D). These results indicate a correlation between endolysosomal membrane cholesterol levels, endolysosomal positioning and dynein clustering.

### Membrane cholesterol levels influence dynein copy number within nano-clusters on endolysosomes

To better understand the mechanisms behind dynein clustering and to causally relate dynein clustering to endolysosomal membrane cholesterol levels, we manipulated cholesterol levels with two commonly used drugs: U18666A that increases endolysosomal membrane cholesterol and Lovastatin that decreases cellular and endolysosomal membrane cholesterol levels (Keyomarsi, 1996; Rocha et al., 2009). ORP1L WT-mCherry-positive endolysosomes in ORP1L KO HeLa cells treated with U18666A indeed had higher membrane cholesterol levels (mean intensity = 3020±183.5 a.u) (by 4.5-fold) compared to those in cells treated with lovastatin (mean intensity = 668.2±47.5 a.u) as measured by filipin intensity (Figure S2A). We will refer to the endolysosomal membrane cholesterol content of the U18666A and Lovastatin treated cells as ‘high cholesterol’ and ‘low cholesterol’ condition, respectively.

Super-resolution imaging revealed that for those endolysosomes that contained dynein nano-clusters associated to them, the number of dynein nano-clusters per endolysosome was independent of cholesterol levels (Figure S2B) but dynein nano-clusters on endolysosomal compartments contained a significantly higher number of localizations under high cholesterol compared to low cholesterol conditions (Figure 2A-B, mean = 71±1.9 for high and 50±1.9 for low cholesterol).

**Figure 2:**
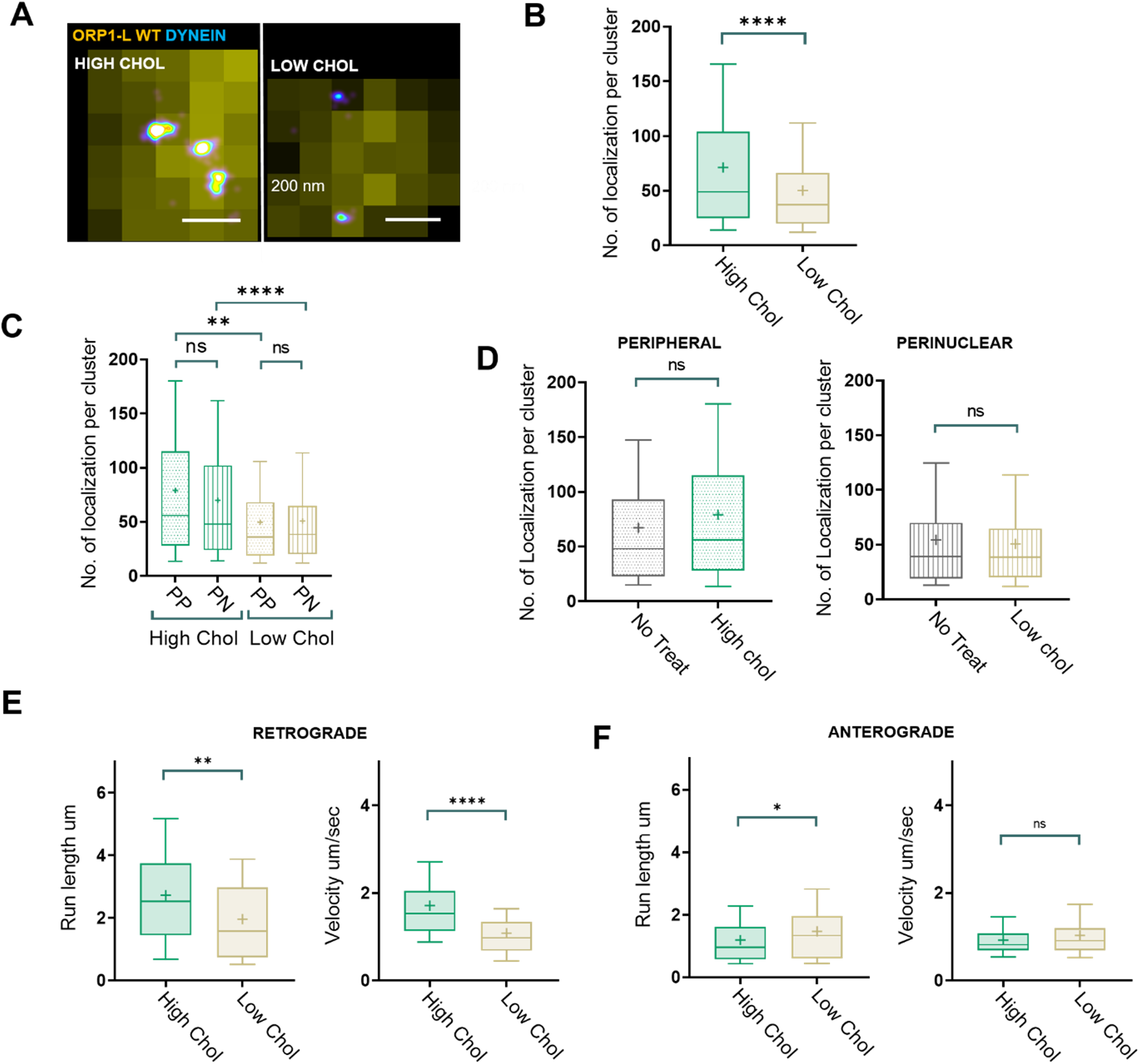
Dynein forms larger nano-clusters containing a higher proportion of dynein multimers on endolysosomes that have high cholesterol content compared to those that have low cholesterol content: (A) An overlay of cropped wide-field image of ORP1L WT-mCherry (yellow) and super-resolution image of immunostained dynein (image is color coded according to localization density with higher density corresponding to white and lower density corresponding to cyan) for an endolysosome treated with U18666A (High Chol) or Lovastatin (Low Chol) in Hela ORP1L-KO cells. *(B) Box plot showing the number of localizations per dynein nano-cluster for endolysosomes in ORP1L WT-mCherry expressing Hela ORP1L-KO cells treated with U18666A (High Chol) (n=224 endolysosomes from n=6 cells, n=2 experiments, mean 71±1.9) versus Lovastatin (Low Chol) (n=217 endolysosomes from n=6 cells, n=2 experiments, mean* 50±1.9*). The box corresponds to 25-75 percentile, the line corresponds to the median, the cross corresponds to the mean and the whiskers correspond to 10-90 percentile. Statistical significance was assessed using a Kolmogorov-Smirnov-test with a p-value <0.0001*. *(C) Box plot showing the number of localizations per dynein nano-cluster for peripherally or peri-nuclearly positioned endolysosomes in ORP1L WT-mCherry expressing Hela ORP1L-KO* cells treated with *U18666A (High Chol PP: n=70 endolysosomes from n=6 cells, n=2 experiments, mean 79±5 and High Chol PN: n=154 endolysosomes from n=6 cells, n=2 experiments, mean 70±2) or Lovastatin (Low Chol PP; n=110endolysosomes from n=6 cells, n=2 experiments, mean 50±2.7 and Low Chol PN; n=107 endolysosomes from n=6 cells, n=2 experiments, mean 51±2.8, respectively). The box corresponds to 25-75 percentile, the line corresponds to the median, the cross corresponds to the mean and the whiskers correspond to 10-90 percentile. Statistical significance was assessed using a Kolmogorov-Smirnov-test with a p-value of n.s.: 0.27, n.s.: 0.93, **: 0.001, ****: <0.0001*. *(D) Box plot showing the number of localizations per dynein nano-cluster for peripherally positioned endolysosomes in ORP1L WT-mCherry expressing Hela ORP1L-KO* cells, *untreated (No Treat) (n=195 endolysosomes from n=6 cells, n=2 experiments, mean 67*±1.9*) and treated with U18666A (High Chol) (n=70 endolysosomes from n=6 cells, n=2 experiments, mean 79*±5*); as well as in peri-nuclearly positioned endolysosomes in untreated cells (No Treat) (n=210 endolysosomes from n=6 cells, n=2 experiments, mean 54*±1.8*) and Lovastatin treated cells (Low Chol) (n=107 endolysosomes from n=6 cells, n=2 experiments, mean 51±2.8). The box corresponds to 25-75 percentile, the line corresponds to the median, the cross corresponds to the mean and the whiskers correspond to 10-90 percentile. Statistical significance was assessed using a Kolmogorov-Smirnov-test with a p-value of 0.1353 and 0.7*. *(E) Box plot showing the retrograde run length of endolysosomes in ORP1L WT-mCherry expressing Hela ORP1L-KO cells* treated with *U18666A (High Chol) (n=135 endolysosomes from n=40 cells, n=3 experiments, mean 2.7±2) or Lovastatin (Low Chol) (n=100 endolysosomes from n=40 cells, n=3 experiments, mean 1.9±1); as well as retrograde velocity of endolysosmes in U18666A treated cells (High Chol) (n=135 endolysosomes from n=40 cells, n=3 experiments, mean 1.7±0.9) or Lovastatin treated cells (Low Chol) (n=100 endolysosomes from n=40 cells, n=3 experiments, mean 1±0.7)*. The box corresponds to 25-75 percentile, the line corresponds to the median, the cross corresponds to the mean and the whiskers correspond to 10-90 percentile. Statistical significance was assessed using a Kolmogorov-Smirnov-test with a p-value of 0.01 and <0.0001. *(F) Box plot showing the anterograde run length of endolysosomes in ORP1L WT-mCherry expressing Hela ORP1L-KO cells* treated with *U18666A (High Chol) (n=135 endolysosomes from n=40 cells, n=3 experiments, mean 1.2±0.8) or Lovastatin (Low Chol) (n=130 endolysosomes from n=40 cells, n=3 experiments, mean 1.4±0.9); as well as anterograde velocity of endolysosmes in U18666A treated cells (High chol) (n=135 endolysosomes from n=40 cells, n=3 experiments, mean 0.9±0.4) versus Lovastatin treated cell (Low Chol) (n=130 endolysosomes from n=40 cells, n=3 experiments, mean 1±0.6). The box corresponds to 25-75 percentile, the line corresponds to the median, the cross corresponds to the mean and the whiskers correspond to 10-90 percentile. Statistical significance was assessed using a Kolmogorov-Smirnov-test with a p-value of 0.01 and 0.17*

These differences were not due to an increase in dynein expression level upon high cholesterol drug treatment, as western blot analysis showed that the level of dynein expression did not increase under high cholesterol treatment (Figure S2C). We once again fit the number of localizations per nano-cluster distribution to the calibration data to determine dynein copy number within nano-clusters under high and low cholesterol conditions, stopping the fit when the residuals stopped changing substantially, which corresponded once again to *f3* (Figure S2D). Once again, we assigned the weight for N=1 IC74 to single dynein (61% and 77% on high and low cholesterol endolysosomes, respectively) and N=3 to multiple (likely 2) copies of dynein (38% and 18% on high and low cholesterol endolysosomes, respectively). The proportion of nano-clusters corresponding to a weight of N=2 IC74s was once again negligible (0.7% and 4% on high and low cholesterol endolysosomes, respectively). This analysis showed that under high cholesterol the proportion of nano-clusters containing two dyneins increased by ~2-fold (from 18% to 38%) (Figure S2D). Finally, low cholesterol levels also led to an increase in the percentage of endolysosomes that completely lacked dynein (Figure S2E, Low chol: 55%, High Chol: 27%).

We next analyzed the cholesterol levels and dynein nano-cluster size on peripheral and peri-nuclear endolysosomes under the two cholesterol treatment conditions. In contrast to physiological conditions, peripheral and peri-nuclear endolysosomes had similar cholesterol levels (Figure S2F, High Cholesterol: mean intensity = 2998±209 a.u. and 3152±180 a.u. for peripheral and peri-nuclear respectively and Low Cholesterol: mean intensity = 491±55 a.u. and 480±43 a.u. for peripheral and peri-nuclear, respectively) as measured by filipin intensity and similar dynein clustering under both high and low cholesterol conditions (Figure 2C, High cholesterol: mean localizations per cluster = 79±9 and 70±2 for peripheral and peri-nuclear endolysosomes, respectively and Low Cholesterol: mean localizations per cluster = 50±2.7 and 51±2.8 for peripheral and peri-nuclear endolysosomes, respectively). Interestingly, the dynein clustering level of peri-nuclear endolysosomes under physiological conditions (mean localizations per cluster = 54±1.8) was similar to those under low cholesterol conditions (mean localizations per cluster = 51±1.8) and the dynein clustering level of peripheral endolysosomes under physiological conditions (mean localizations per cluster = 67±1.9) was similar to those under high cholesterol conditions (mean localizations per cluster = 79±5) (Figure 2D). These results indicate that cholesterol levels and not endolysosome positioning determine the level of dynein clustering on endolysosomal membranes.

To determine whether these differences in dynein clustering correlated with changes in endolysosomal transport, we determined the positioning and transport properties of endolysosomes under high and low cholesterol conditions. It has previously been shown that high cholesterol leads to peri-nuclear clustering of endolysosomes whereas low cholesterol leads to their peripheral scattering (Rocha et al., 2009). Here we further confirmed that the positioning of endolysosomes was indeed impacted by cholesterol levels, with low cholesterol leading to more scattered and high cholesterol leading to more peri-nuclearly clustered endolysosomes (Figure S2G), consistent with previous results. In addition, we measured the retrograde and anterograde run length and the velocity of endolysosomes in live cell movies under low and high cholesterol conditions (Figure 2E and F). While high cholesterol led to increased run length and velocity as compared to low cholesterol in the retrograde direction (Figure 2E, mean run length = 2.7±2 μm and 1.9±1 μm for high and low cholesterol and mean velocity = 1.7±1 μm/sec and 1±1 μm/sec for high and low cholesterol, respectively), the anterograde run length was only slightly affected (slightly longer run length in low cholesterol as compared to high cholesterol) and the anterograde velocity was unchanged by cholesterol treatment (Figure 2F, mean run length = 1.2±0.8 μm and 1.4±0.9 μm for high and low cholesterol and mean velocity = 0.9±0.4 μm/sec and 1±0.6 μm/sec for high and low cholesterol, respectively). These results further suggest that increased dynein clustering under high cholesterol likely leads to more efficient retrograde transport whereas kinesin dependent anterograde transport is not affected.

Taken together our results show that membrane cholesterol levels impact not only dynein recruitment but importantly also the level of dynein clustering and the proportion of dynein multimers within nanoclusters on endolysosomal membranes.

### The cholesterol sensing domain of ORP1L regulates ORP1L clustering on endolysosomal membranes in a cholesterol dependent manner

Dynein does not directly bind to endolysosomal membranes and the mechanisms that can lead to the formation of nano-clusters with increased dynein copy number are unknown. Dynein is recruited to endolysosomes through a tripartite complex of Rab7-ORP1L-RILP (Johansson et al., 2007; Pfeffer, 2001; Rocha et al., 2009). ORP1L contains multiple lipid binding domains that allow it to bind to either oxysterols or phospholipids (Johansson et al., 2005; Olkkonen and Li, 2013; Zhao and Ridgway, 2017). Hence, we asked whether the cholesterol sensing domain of ORP1L is responsible for regulating dynein clustering on endolysosomal membranes. To address this question, we first imaged ORP1L’s spatial distribution on endolysosomes in wild type or ORP1L-KO HeLa cells that express ORP1L WT-mCherry (Zhao and Ridgway, 2017) under low or high cholesterol treatment conditions using super-resolution microscopy (Figure 3A). We again analyzed isolated, round ORP1L WT-mCherry positive endolysosomes within a size range of 50-700 nm in the super-resolution images. ORP1L appeared uniformly distributed (i.e. a single, large ORP1L cluster covering the entire endolysosomal membrane) under low cholesterol conditions and more clustered (i.e. multiple, smaller clusters covering only part of the endolysosomal membrane) under high cholesterol conditions (Figure 3A).

**Figure 3:**
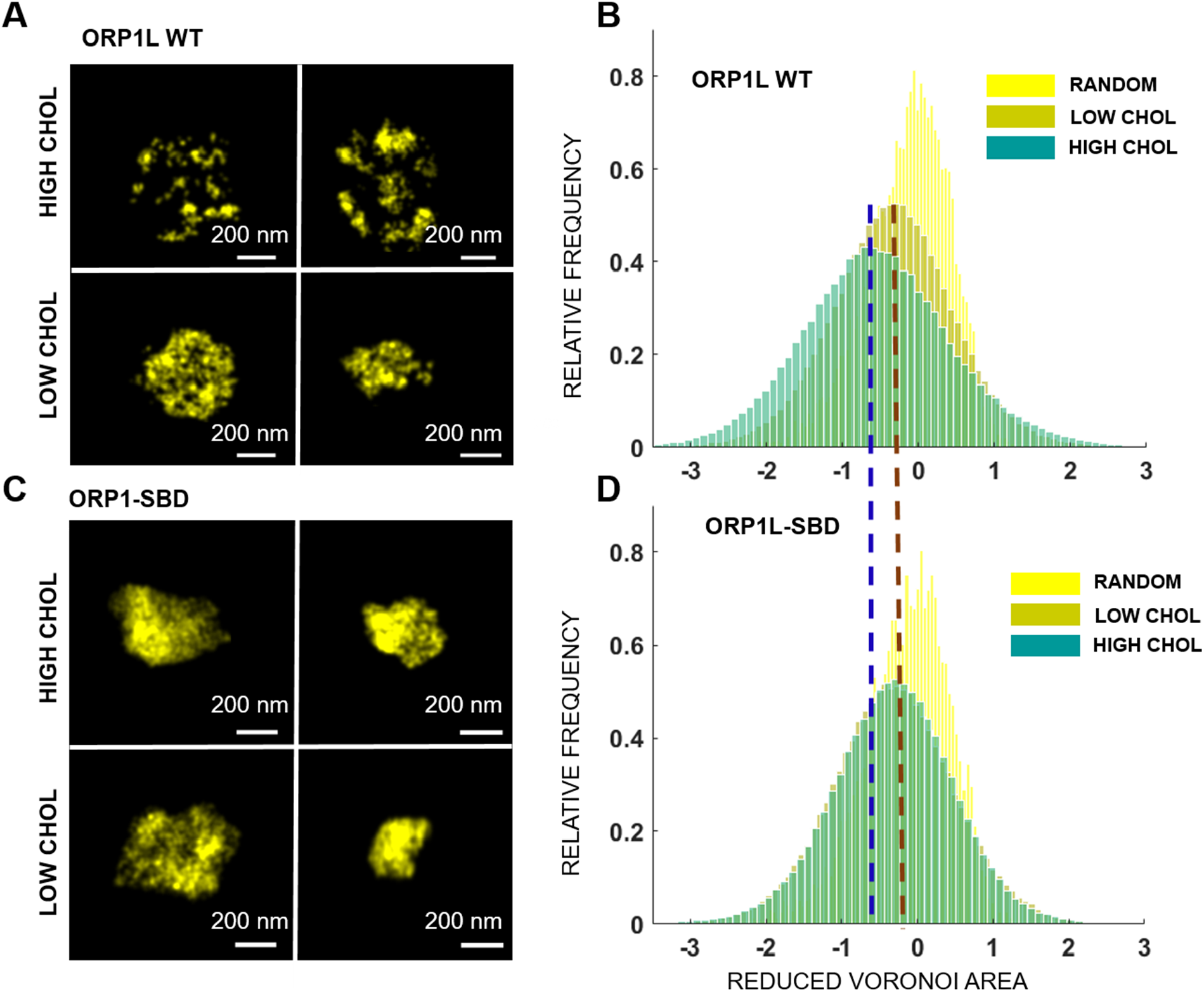
ORP1L is more clustered on endolysosomes having higher cholesterol levels in a manner dependent on its cholesterol binding domain: (A) Super-resolution images of full length ORP1L (ORP1L WT-mCherry) in Hela ORP1L-KO cells treated with U18666A (High Chol, upper panels) and Lovastatin (Low Chol, lower panels). (B) Log plot of the Reduced Voronoi Polygon area distribution for super-resolution images of full length ORP1L (ORP1L WT-mCherry) in Hela ORP1L-KO cells treated with U18666A (dark green, High Chol) (n=95 endolysosomes from n=6 cells, n=2 experiments), Lovastatin (light green, Low Chol) (n=107 endolysosomes from n=6 cells, n=2 experiments) and for a random distribution of localizations (yellow, Random). The dashed lines are a guide to the eye to highlight the shift in the peak position of Reduced Voronoi Polygon area distributions with cholesterol treatment for full length ORP1L (ORP1L WT-mCherry) and sterol binding deficient ORP1L mutant lacking residues 560-563 (ORP1L SBD-mCherry). (C) Super-resolution images of sterol binding deficient ORP1L mutant lacking residues 560563 (ORPiL SBD-mCherry) in Hela ORP1L-KO cells treated with U18666A (High Chol, upper panels) and Lovastatin (Low Chol, lower panels). (D) Log plot of the Reduced Voronoi Polygon area distribution for super-resolution images of sterol binding deficient ORPiL mutant lacking residues 560-563 (mCherry-ORP1L-SBD) in Hela ORP1L-KO cells treated with U18666A (dark green, High Chol) (n=120 endolysosomes from n=6 cells, n=2 experiments), Lovastatin (light green, Low Chol) (n=170 endolysosomes from n=6 cells, n=2 experiments) and for a random distribution of localizations (yellow, Random).

To quantify the level of ORP1L clustering, we once again used Voronoi tessellation to segment individual ORP1L clusters on endolysosomal membranes. This approach revealed an increased number of ORP1L clusters with smaller area under high cholesterol (Figure S3A, mean clusters per endolysosome = 2.6±0.32 with a mean area of 23×10^3^ nm^2^) compared to low cholesterol (mean clusters per endolysosome = 1±0.04 with a mean area of 10×10^4^ nm^2^) conditions. However, we also found that, surprisingly, the localization density of ORP1L was higher on endolysosomes under low cholesterol compared to high cholesterol conditions (Figure S3B, mean localizations per area = 0.017±0.0006 for low and 0.007±0.0003 for high cholesterol), suggesting that more ORP1L binds to endolysosomes when their membrane cholesterol levels are lower. The localization density of ORP1L under high or low cholesterol was consistent between three independent biological replicates and the high cholesterol treatment consistently had lower localization density compared to the low cholesterol treatment, suggesting that ORP1L expression levels were consistent among experimental conditions (Figure S3C). When endogenous ORP1L clustering level was quantified on CD63 positive endolysosomal compartments in wild type Hela cells using an ORP1L antibody, it also was more clustered under high cholesterol (mean clusters per endolysosome = 4±0.31 and 2±0.02, with a mean area of 14×10^3^ nm^2^ and 45×10^3^ nm^2^ under high and low cholesterol, respectively) and had lower localization density (mean localizations per area = 0.005±0.0002 and 0.008±.0003 under high and low cholesterol, respectively) under high cholesterol compared to low cholesterol conditions (Figure S3D-E), further demonstrating that the results are not an artifact of ORP1L over-expression.

To ensure that the increased protein density with low cholesterol treatment does not confound the clustering analysis using Voronoi tessellation, which is dependent on localization density, we developed an alternative quantification method that is insensitive to differences in localization density and protein amount (see Methods). To this end, we carried out Voronoi tessellation and re-scaled the distribution of Voronoi polygon areas so that the mean area was set to unity such that we could measure the clustering tendency of ORP1L without explicitly compensating for different localization densities. For a clustered distribution, we expected to see a shift in the mode of the reduced Voronoi polygon area distribution towards a smaller value. Indeed, the mode of the distribution for endolysosomal compartments under high cholesterol conditions was shifted to smaller polygon areas compared to low cholesterol conditions or compared to a simulated random distribution, indicating that ORP1L is more clustered on endolysosomes with high membrane cholesterol levels (Figure 3B). We further used a statistical test (Kullback-Leibler Divergence or KL Divergence, see Methods) to determine a clustering tendency score that shows how much ORP1L organization differed from a random distribution. The clustering tendency score was 0.75 under high cholesterol and 0.14 under low cholesterol (a 5-fold difference). These results further confirm that ORP1L’s organization on endolysosomal membranes deviated significantly more from a random distribution under high cholesterol compared to low cholesterol conditions. Overall, these super-resolution data show that ORP1L, like dynein, forms a similar number of nano-clusters on endolysosomes having higher membrane cholesterol levels.

To determine if the differences in ORP1L’s spatial distribution on endolysosomes were due to its cholesterol binding, we expressed mCherry fused to a sterol binding deficient ORP1L mutant lacking the residues 560–563 (ORP1L SBD-mCherry) (Vihervaara et al., 2011; Zhao and Ridgway, 2017) in HeLa ORP1L KO cells (Figure 3C-D and Figure S3F-G). Super-resolution images of the ORP1L SBD-mCherry mutant (Figure 3C) and both Voronoi cluster segmentation analysis (Figure S3F, mean clusters per endolysosome = 1±.03 for high and 1.1±.04 for low cholesterol conditions) and reduced Voronoi polygon area distribution analysis (Figure 3D, Clustering tendency score for ORP1L-SBD: 0.1 for high and 0.09 for low cholesterol conditions) showed that the distribution of the ORP1L-SBD mutant was uniform on endolysosomal membranes independent of cholesterol levels. The membrane localization density of this mutant was overall high under both high and low cholesterol conditions (Figure S3G, mean localizations per area = 0.034±.0017 and 0.026±.00075 for high and low cholesterol, respectively) and at a similar level to the full length ORP1L under low cholesterol conditions (Figure S3B, mean localizations per area = 0.017±0.00062). These results suggest that under low cholesterol or when ORP1L lacks its cholesterol sensing domain, it is recruited to and binds phospholipids on endolysosomes at a high level and likely in a non-specific manner.

Overall, these results strongly support that the cholesterol sensing domain of ORP1L regulates its spatial distribution on endolysosomal membranes in a cholesterol dependent manner, with high cholesterol leading to more specific ORP1L recruitment (Figure S3B) and formation of ORP1L nano-clusters (Figure 3A-B).

### The cholesterol sensing domain of ORP1L regulates dynein clustering and endolysosomal positioning in a cholesterol dependent manner

Having established that ORP1L’s cholesterol sensing domain regulates ORP1L’s level of clustering on endolysosomal membranes in a cholesterol dependent manner, we next asked if ORP1L’s cholesterol sensing domain also impacts dynein clustering. We thus imaged dynein using super-resolution microscopy in ORP1L-KO HeLa cells expressing the ORP1L SBD-mCherry mutant. Analysis of dynein nano-clusters on endolysosomes under high and low cholesterol conditions showed that dynein clustering was no longer sensitive to cholesterol levels in cells expressing the ORP1L SBD-mCherry mutant as the sole ORP1L isoform (Figure 4A-B, mean localizations per cluster = 51±1.2 for high and 53±0.88 for low cholesterol). In addition, the level of dynein clustering in cells expressing the ORP1L SBD-mCherry mutant was similar to the level of dynein clustering under low cholesterol conditions in cells expressing the full length ORP1L WT-mCherry (mean localizations per cluster = 50±1.9) (compare Figure 2B and Figure 4B).

**Figure 4:**
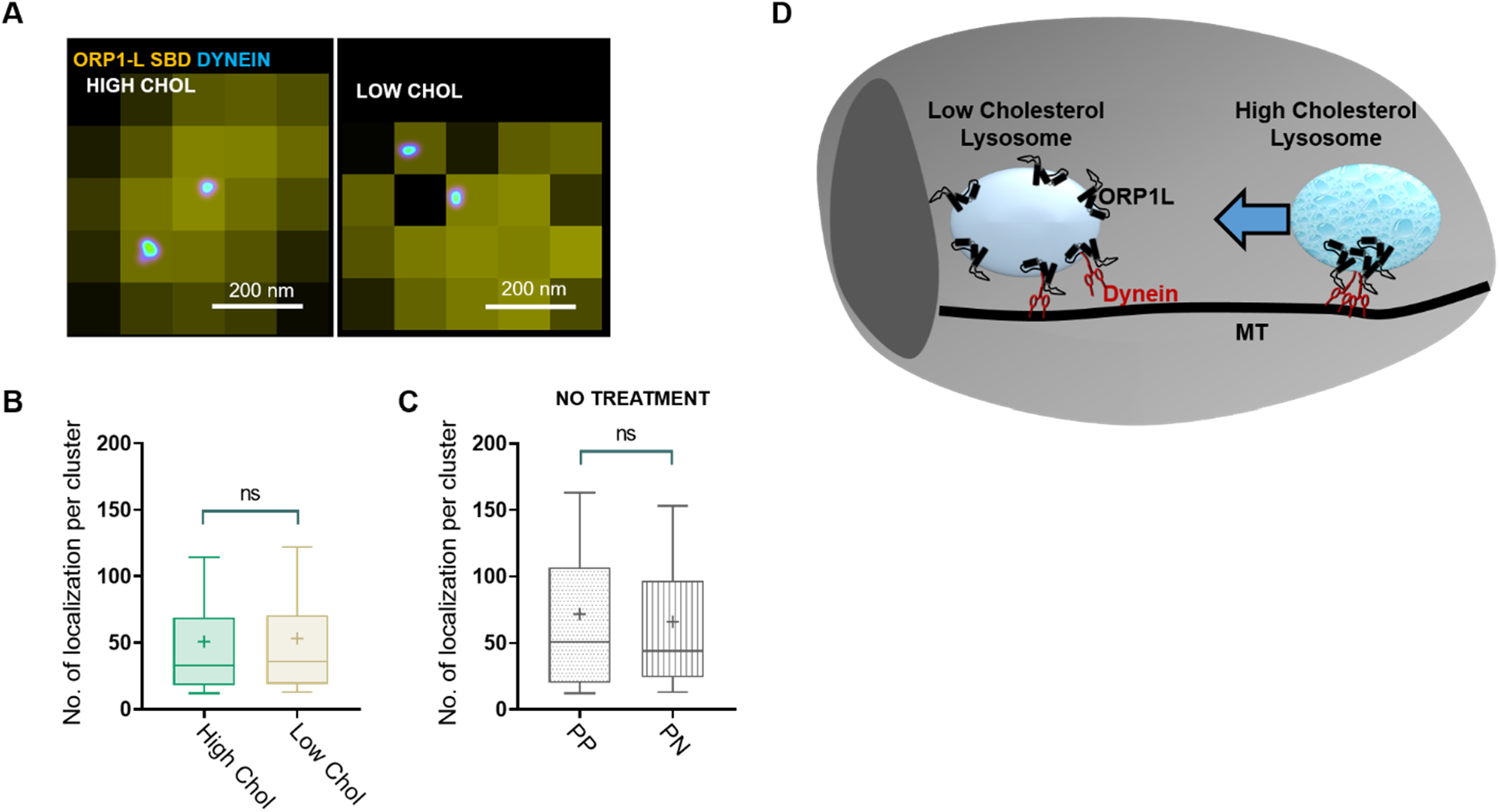
Dynein nano-clusters are insensitive to cholesterol levels in cells expressing sterol binding deficient ORP1L mutant: (A) An overlay of cropped wide-field image of ORP1L WT-mCherry (yellow) and super-resolution image of immunostained dynein (image is color coded according to localization density with higher density corresponding to white and lower density corresponding to cyan) in Hela ORP1L-KO cells expressing sterol binding deficient ORP1L mutant lacking residues 560-563 (ORP1L SBD-mCherry) treated with U18666A (High Chol) or Lovastatin (Low Chol) *(B) Box plot showing the number of localizations per dynein nano-cluster for endolysosomes in Hela ORP1L-KO cells expressing the sterol binding deficient ORP1L mutant lacking residues 560-563 (ORP1L SBD-mCherry) and treated with U18666A (High Chol) (n=450 endolysosomes from n=6 cells, n=2 experiments, mean 51*±1.2*) or Lovastatin (Low Chol) (n=300 endolysosomes from n=6 cells, n=2 experiments, mean 53±0.88). The box corresponds to 25-75 percentile, the line corresponds to the median, the cross corresponds to the mean and the whiskers correspond to 10-90 percentile. Statistical significance was assessed using a Kolmogorov-Smirnov-test with a p-value of 0.06*. (C) Box plot showing the number of localizations per dynein nano-cluster for peripherally positioned endolysosomes (PP) (n=100 endolysosomes from n=5 cells, n=2 experiments, mean 71±6) and peri-nuclearly positioned endolysosomes (PN) (n=150 endolysosomes from n=5 cells, n=2 experiments, mean 66±2) in Hela ORP1L-KO cells expressing the sterol binding deficient ORP1L mutant lacking residues 560-563 (ORP1L SBD-mCherry). The box corresponds to 25-75 percentile, the line corresponds to the median, the cross corresponds to the mean and the whiskers correspond to 10-90 percentile. Statistical significance was assessed using a Kolmogorov-Smirnov-test with a p-value of 0.44. (D) Proposed model of how cholesterol levels and ORP1L regulate dynein nano-clustering and endolysosome positioning: Peripheral lysosomes with higher cholesterol content have a more clustered organization of ORP1L and dynein likely leading to efficient retrograde transport (blue arrow). Perinuclear lysosomes having lower cholesterol content have more uniform ORP1L organization leading to lower dynein recruitment and clustering likely facilitating endolysosome anchoring at the peri-nuclear region or anterograde trafficking back to the cell periphery.

While in cells expressing the ORP1L WT-mCherry, peripheral endolysosomes had more clustered ORP1L (mean clusters per endolysosome = 1.4±0.7) compared to peri-nuclear endolysosomes (mean clusters per endolysosome = 1±0.2), these differences disappeared in cells expressing the ORP1L SBD-mCherry mutant (mean clusters per endolysosome = 1.1±0.4 and 1.1±0.6 for peripheral and peri-nuclear, respectively) (FigureS4A-B). The sub-cellular positioning of endolysosomes also became insensitive to cholesterol treatment in cells expressing the ORP1L SBD-mCherry mutant and the endolysosomes were overall more scattered throughout the cell under high or low cholesterol conditions as well as under physiological conditions (Figure S4C). Finally, peripheral and peri-nuclear endolysosomes had similar level of dynein clustering in untreated cells expressing the ORP1L SBD-mCherry mutant (Figure 4C, mean number of localizations = 71±6 for high and 66±2 for low cholesterol). Taken together, these results suggest that the higher copy number of dynein within nano-clusters on peripherally located endolysosomes with higher cholesterol content is dependent on the cholesterol binding domain of ORP1L (Figure 4D).

## Discussion

Here using super resolution microscopy and quantitative analysis, we visualize the spatial organization of dynein motor and its adapter protein ORP1L on endolysosomal membranes. We find that dynein forms nano-clusters consisting of mainly single but also a small but significant proportion of multiple copies of dynein motor. We note that the proportion of dynein multimers are likely underestimated in our quantitative analysis due to steric effects impacting labeling efficiency. Using perturbation experiments in which we manipulated endolysosomal membrane cholesterol levels we show that the copy number of dynein within nano-clusters on endolysosomal membrane is increased under high cholesterol. This increased clustering is regulated by the cholesterol binding ability of ORP1L that forms part of the Rab7-ORP1L-RILP tripartite complex, which recruits dynein to endolysosomal membranes.

Cryo-EM experiments demonstrated that BICD2 and Hook3 bind two copies of dynein motors (Urnavicius et al., 2018) but whether the endolysosomal adapter proteins Rab7-ORP1L-RILP also recruit multiple dyneins is unknown. Our results demonstrate that besides the stoichiometry between motor proteins and their adapter proteins, additional mechanisms inside cells are likely at play to increase motor protein clustering. We show that cholesterol levels and the ability of the adapter protein ORP1L to sense and bind cholesterol impact the copy number of dynein motor within nano-clusters on endolysosomal membranes. High cholesterol treatment increased the proportion of nano-clusters containing dynein multimers. Peripheral endolysosomes also had a higher proportion of nano-clusters with 2 copies of dynein compared to perinuclear endolysosomes. The number of dynein nano-clusters per endolysosome, on the other hand, was independent of cholesterol or endolysosome position. These results suggest that under high cholesterol, more dynein is recruited into the dynein nano-clusters. An interesting hypothesis that future experiments can test is that there are pre-existing hot spots, potentially cholesterol enriched domains containing ORP1L, which may act as recruitment sites for multiple copies of dynein.

Endolysosome run length and velocity were also higher in the retrograde direction under high cholesterol whereas anterograde run length and velocity was cholesterol independent. These results suggest that having dynein present in multiple copies may be important for efficient retrograde transport, which is consistent with the recent Cryo-EM and in vitro single molecule imaging data showing that coupled dyneins move more processively and faster than single dynein (Urnavicius et al., 2018).

Interestingly, ORP1L was present in higher amounts on endolysosomes under low cholesterol compared to high cholesterol and its spatial distribution was uniform rather than clustered under low cholesterol. However, increased amounts of ORP1L did not lead to more dynein recruitment, to the contrary, the number of dynein motors within nano-clusters as well as the total number of dynein motors on endolysosomes were decreased under low cholesterol. This result may be due to the fact that ORP1L needs to be in a tripartite complex with Rab7 and RILP to recruit dynein and such complex formation maybe compromised under low cholesterol conditions in which ORP1L may bind the endolysosome membrane in a more non-specific manner. Future experiments probing the spatial co-relationship between ORP1L, Rab7, RILP and dynein can address this possibility.

Previous *in vitro* work showed that early phagosomes engulfing 2-micron sized polystyrene beads isolated from dictyostelium cells were completely uniformly covered with dynein on their membrane whereas late phagosomes that move unidirectionally towards the retrograde direction had a highly clustered dynein distribution (Rai et al., 2016). These changes in dynein distribution were correlated to the membrane cholesterol content of phagosomes. In contrast, our results show that instead of uniformly covering the entire membrane, only a small number of dyneins are associated to endolysosomal membranes *in vivo*. A dramatic shift from a uniform dynein coverage to a highly clustered dynein distribution hence is likely not needed for regulating retrograde transport of native cellular compartments as these previous experiments with non-native compartments suggested. Instead, a small shift from mainly single copies of dynein to a small proportion of dynein mutimers is likely sufficient to bias retrograde transport of native sub-cellular compartments. Our results are more in line with high speed dark field microscopy and optical trapping experiments on axonal endosomes, which also suggested that up to 8 dynein motors can dynamically cluster under load (Chowdary et al., 2018).

Interestingly, the number of dyneins within nano-clusters is higher for peripherally located endolysosomes, which also have higher cholesterol levels. These results may seem counterintuitive as dynein is responsible for retrograde transport and hence one may expect more dynein clustering on endolysosomes accumulating in the peri-nuclear region. However, once endolysosomes reach the peri-nuclear region, they may no longer require clustered dynein multimers to maintain their peri-nuclear positioning. Dynein may dissociate from the membrane to be recycled once endolysosomes are positioned in the peri-nuclear region. On the other hand, peripherally located endolysosomes may depend on clustered dynein multimers to be transported to and deposited at the peri-nuclear region. In addition, it is known that endolysosomes contact the endoplasmic reticulum as they are trafficked within the cell cytoplasm (Friedman et al., 2013; Rocha et al., 2009; Zhao and Ridgway, 2017). Such contacts lead to exchange of membrane lipids and maturation of endolysosomes (van der Kant et al., 2013; Zhao and Ridgway, 2017). It is plausible that multiple endolysosome-ER contacts during retrograde transport play a role in lowering the cholesterol content of endolysosomes, potentially providing a mechanism for dissociation of dynein and decreased dynein clustering once the endolysosomes reach their peri-nuclear destination. Future correlative live-cell and super-resolution imaging experiments (Balint et al., 2013; Mohan et al., 2019; Verdeny-Vilanova et al., 2017) will enable directly linking the transport properties of endolysosomes to the level of dynein clustering on their membrane.

Previous studies showed that ORP1L, depending on its conformation, can either bind dynein or make contacts with the ER-membrane (Olkkonen and Li, 2013; Rocha et al., 2009; Wijdeven et al., 2016; Zhao and Ridgway, 2017). A change in ORP1L’s conformation leads to shedding of dynein and initiation of contact between ORP1L and the ER-membrane protein VAP to regulate endolysosomal positioning (Rocha et al., 2009). Our results are consistent with these former studies as we show that in addition to the level of dynein clustering, the recruitment of dynein to endolysosomal compartments is also dependent on cholesterol levels. Here, we additionally show that ORP1L forms nano-clusters on endolysosome membranes when the membrane cholesterol level is high. In addition, not only more dynein is recruited to the endolysosomal membrane under these conditions but also the proportion of dynein present as multimers within nano-clusters is increased. Hence, multiple mechanisms, including recruitment of more dynein motors and clustering of the recruited dynein motors on the endolysosomal membrane, may be at play to increase the efficiency of retrograde transport and regulate endolysosomal positioning. It will be interesting in the future to examine the differential impact of having more dynein that is not present as multimers within nano-clusters versus having similar amount of dynein clustered in multimers within nano-clusters on the transport and positioning of endolysosomes. It is plausible to hypothesize that increased dynein recruitment in the absence of dynein clustering is not sufficient for enhancing retrograde transport.

Overall, we propose an *in vivo* model dependent on cholesterol levels by which multiple dyneins can be recruited and clustered on endolysosomal membranes leading to their efficient retrograde transport and positioning (Fig. 4D). Increased dynein clustering in response to cholesterol levels is likely to be functionally significant as it impacts the sub-cellular positioning of endolysosomal compartments. Metabolic disorders that lead to accumulation of lipids including cholesterol in endolysosomal compartments like Niemann Pick Disease (NPC) are typically associated with alterations in endolysosomal homeostasis and function (Torres et al., 2017). In the future it would be interesting to explore if the nanoscale organization of ORP1L and dynein is altered on endolysosomal membranes in NPC and other lysosomal storage disorders leading to their mislocalization within the cell and whether restoring the proper nanoscale organization of these cytoskeletal proteins can restore endolysosomal function. It would also be interesting to determine if similar mechanisms play a role in regulating kinesin clustering or in regulating transport of other organelles including Golgi vesicles and autophagosomes. It would further be exciting to determine the precise stoichiometry of adapter-motor complexes on organelle membranes to determine how the stoichiometry can be precisely tuned to regulate organelle transport and positioning. Our work establishes the methodology needed and opens the door to carry out these future studies.

## Materials and Methods

### Cells and transfections

Wild type HeLa cells were obtained from the American Type Culture Collection (CCL-2, ATCC, Manassas, VA). HeLa-ORP1L-null cell lines, as well as mCherry-tagged ORP1L and ORP1L-SBD constructs were a kind gift from Prof. Neale Ridgway (Dalhousie University, Depts. of Pediatrics, and Biochemistry and Molecular Biology, Atlantic Research Centre, Halifax, Nova Scotia, Canada). HeLa cells were grown in DMEM (GIBCO Laboratories, Grand Island, NY) supplemented with 10% fetal bovine serum and antibiotics, and maintained in 5% CO2 at 37°C. Cells were transiently transfected with mCherry-tagged wild type or mutant ORP1L at 70% confluency using Lipofectamine 2000 reagent (Invitrogen) according to the manufacturer’s protocol. Cells were subjected to experimental treatments 24 h after transfection.

### Pharmacological treatment of cells

Lovastatin (Sigma 1370600) was converted from its inactive prodrug form to its active open acid form by dissolving Lovastatin (52gms) in ethanol (95%, 1.04 ml), followed by addition of 1N NaOH (813 μl), followed by heating for 2 hrs at 50°C and neutralized with 1N HCL (pH 7.2). The volume was made up to 13 ml by adding distilled water, giving 10mM active Lovastatin solution later aliquoted and stored at −20°C(Keyomarsi, 1996). Mevalonic acid lactone (Sigma M4667, 1 gm) was converted to its active form by dissolving in ethanol (3.5 ml), followed by addition of 1N NaOH (4.2 ml) and heating for 2 hrs at 55°C. The solution was made up to 15.4 ml with distilled water and neutralized with 1N HCL (pH 7.2), giving 500 mM of stock solution, later aliquoted and stored at −20. U18666A (Sigma U3633) was dissolved in ethanol giving final concentration of 10mg/ml(Rocha et al., 2009). For cholesterol depletion treatment, cells were cultured in DMEM, 10% Lipoprotein deficient serum (Sigma S5394), 50uM Lovastatin, 230uM Mevalonate for 6 hrs before fixing for immunostaining. For high cholesterol treatment cells were cultured in DMEM, 10% FBS and 3μg/ml U18666A for 12 hrs before fixing.

### Immunostating

Hela ORP1L KO cells were transiently transfected with ORP1L WT-mCherry or ORP1L SBD-mCherry, treated with U18666A/Lovastatin or not treated and immunostained for ORP1L and dynein. Endogenous ORP1L was immunostained in untransfected Hela cells treated with U18666A/Lovastatin or not treated. For ORP1LWT/SBD-mCherry and endogenous ORP1L immunostaning, cells were fixed with 4% (Vol/Vol) Paraformaldehyde in PBS for 20 mins and for dynein immunostaining, cells were fixed in prechilled 1:1 Ethanol/Methanol solution for 3 mins on ice. The cells were blocked in blocking buffer (3% BSA, 0.2% Triton X-100 in PBS) for 1 hr. Cells were incubated with primary anitbodies: Chicken anti-mCherry (Novus biotech nbp2-2515, 1:500), mouse anti-dynein (Abcam ab23905, 1:50 or 1:5000 for single dynein imaging) and rabbit anti-ORP1L (Abcam ab131165, 1:100) in blocking buffer for 1 hr on a rocker. Cells were washed with washing buffer (0.2% blocking buffer, .05% Triton X-100 in PBS) three times. Custom made secondary antibodies were labeled with an Alexa Fluor 405-Alexa Fluor A647 activator/reporter dye pair combination at concentrations (0.1-0.15 mg/μl) and used in the ratio of 1:50 in blocking buffer for 40 mins, at RT on a rocker. Sample was then washed three times in PBS.

### Filipin staining

Filipin (Sigma F4767) was lyophilized, aliquoted (250 μg per aliquot) and stored at −80°C. Filipin was resuspended in 5 μl DMSO. Cells were fixed in 4% Paraformaldehyde for 20 mins and then rinsed 3 times with PBS. Background autofluorescence was quenched with 50mM NH4CL for 10 mins. Cells were incubated for 2 hrs with 100 μg/ ml working solution of Filipin in 3% BSA. Cells were washed with 1% BSA in PBS three times before imaging. Imaging was performed immediately.

### Western blot

Western blot analysis was performed using the two-color Odyssey LI-COR (Lincoln, NE) technique according to the manufacturer’s protocol. A rabbit monoclonal antibody to ORP1L (ab131165, Abcam), mouse monoclonal antibody to dynein (ab23905, Abcam), and a mouse monoclonal antibody to detect GAPDH (clone 3B1E9, GenScript A01622–40) were used at a dilution of 1:1,000 in blocking buffer. The secondary antibody IRDye800CW Donkey anti-Rabbit and IRDye680RD Donkey anti-Mouse (LI-COR) were used in 1:10000 dilution for imaging in the green 800-nm and red 700-nm channels, respectively.

### STORM Imaging

Single-molecule imaging was done using imaging buffer comprising of 50 mM Tris, pH 7.5, 10 mM NaCl, 0.5 mg/mL glucose oxidase (Sigma, G2133), 40 μg/mL catalase (Roche Applied Science, 106810), 10% (w/v) glucose and 10% (v/v) Ciseamine (77mg/ml of 360mM HCL) (Bates et al., 2007). Images were acquired on the Oxford Nanoimager-S microscope which has the following configuration: 405, 488, 561, and 640 nm lasers, 498–551 and 576–620 nm band-pass filters in channel 1, and 665–705 nm band-pass filters in channel 2, 100× 1.4 NA objective (Olympus), and a Hamamatsu Flash 4 V3 sCMOS camera. Localizations were acquired with 10-ms exposure over 50,000 frames with 405 nm activation and 647 nm excitation. Images were processed and localizations were obtained using the NimOS localization software (Oxford Nanoimaging).

### Live cell imaging

Hela ORP1L KO cells were transfected with ORP1L WT-mCherry and treated with U18666A/Lovastatin for 6 hrs, as most endolysosomes were peri-nuclear after 12 hrs of treatment with U18666A. Videos were acquired on the Oxford Nanoimager-S microscope with 100 ms exposure over 300 frames, 561 nm excitation with HiLo illumination and 37°C sample temperature.

### Data Analysis

#### STORM data Analysis

To identify dynein clusters on ORP1L positive compartments, the intensity profile of conventional ORP1L image was used as a mask. Line intensity profile of the ORP1L conventional image along x and y axis was plotted on image J, and the width across one third of the full intensity maxima was considered as the mask (Figure S1B).

For quantitative analysis we used custom written MATLAB codes. A previously described method was adapted that segments super-resolution images based on Voronoi tessellation of the fluorophore localizations (Andronov et al., 2016; Levet et al., 2015). Voronoi tessellation of a STORM image assigns a Voronoi polygon to each localization, such that the polygon area is inversely proportional to the local localization density. The spatial distribution of dynein or ORP1L localizations from each ORP1L positive endolysosome is represented by a set of Voronoi polygons such that smaller polygon areas correspond to regions of higher density. The Voronoi polygons at the endolysosomal edge are extremely large and were omitted for any quantification. Dynein and ORP1L clusters were segmented by grouping adjacent Voronoi polygons with areas less than a selected threshold and imposing a minimum number of localizations. For ORP1L WT-mCherry, ORP1L SBD-mCherry, dynein (ORP1L positive endolysosomes) and dynein (ORP1L-SBD positive endolysosomes), the selected area thresholds were 0.0156 px^2^, 0.02 px^2^, 0.01 px^2^ and, 0.01 px^2^, respectively and the minimum number of localizations imposed were16,10, 7 and 7, respectively. ORP1L localization density was calculated by normalizing the total number of localizations per endolysosme by endolysosome area. Each endolysosme area was calculated by summing up all of its Voronoi polygon areas. The low cholesterol treatment yielded endolysosomes with ORP1L localization densities 4.5 times higher as compared to the high cholesterol treated endoslysosome localization densities. To compare endoslysomal ORP1L distribution following cholesterol treatments, the ORP1L localization densities should be comparable. To address this we divided the Voronoi polygon areas of each endolysosome by its mean Voronoi polygon area, such that the mean localization density of each endolysosome is in reduced units of 1. All reduced Voronoi polygon areas from each endolysosme for each treatment were pulled together, and the histogram was plotted. The distribution of Voronoi areas from uniformly simulated random points is fit to an analytical distribution (Tanemura, 2003). A method for calculating the KL divergence between two histograms or between a histogram and an analytical distribution (Perez-Cruz, 2008) was implemented in Matlab. The KL divergence scores between the experimental reduced voronoi areas and the theoretical random distribution were calculated to determine the clustering tendency score of each cholesterol treatment. ORP1L reduced Voronoi Area analysis and number of clusters per endolysosome analysis were done with endolysosmes with radius < 350nm. Dynein number of cluster per endolysosome analysis has been done with endolysosmes with radius < 350nm and dynein localizations per cluster analysis has been done with endolysosmes with radius <400 nm.

#### Dynein copy number quantification

For single dynein IC74 calibration analysis (100 fold diluted dynein antibody), dynein localizations were taken from the whole cell except the nucleus region to avoid biased results and Voronoi analysis with subsequent cluster analysis was performed. For quantification of dynein copy numbers, the single dynein data (localizations per cluster) was fit to a lognormal distribution function to obtain the μ and σ values for the calibration function (*f_1_*). Using these values, the peripheral, perinuclear and high, low cholesterol dynein data (localizations per cluster) were fit with functions (*f_1_, f_2_, f_3_*…) with *f_2_, f_3_*. corresponding to linear convolutions of *f_1_* with itself and having weights of *w_1_, w_2_, w_3_*…. Residuals were calculated after each fit (*f_1_* alone corresponding to N=1, or *f_1_* and *f_2_*corresponding to N=2, *f_1_, f_2_* and *f_3_* corresponding to N=3 etc…). The fitting was stopped at N=3 with the criterion that the residuals did not vary after N=3.

#### Fillipin Intensity

Filipin intensity was calculated using Image J ‘plot profile’ tool. All ORP1L positive endolysosomes were analyzed by drawing a line segment across it and looking at its intensity profile using the ‘plot profile’ tool. The average of 3-4 highest intensity points were taken and the average background intensity was subtracted from that.

#### Live cell tracking

ORP1L WT-mCherry positive endolysosomes moving towards nucleus was considered retrograde transport and the ones moving towards cell peripheral and away from the nucleus was considered anterograde transport. Endolysosomal positions were tracked by a semiautomated, custom-written, particle-tracking software. ORP1L mCherry positiveTrajectories were analyzed by performing a 2D Gaussian fit to the point spread function and the *x* and *y* positions were determined.

A previously written custom MATLAB program was used to determine the active and passive phases of the endolysosomal trajectory, as previously described (Verdeny-Vilanova et al., 2017). As previously described and validated (Verdeny-Vilanova et al., 2017), a moving window analysis (four-point segments) was performed along the trajectory data points and the ratio between the total displacement (the initial and final points of the segment) and the sum of displacements between the points within the segment was calculated. This ratio provides an estimate of the linearity of the segment, and values close to 1 were considered active phases. A threshold ratio of 0.6 was considered to distinguish between active and passive phases. Additional criteria like calculating power-law exponent from the mean square displacement (MSD) and angle criterion (successive displacement vectors showing angles less than 90° were categorized as passive) were imposed to confirm the active or passive categorization. For segments categorized as passive but containing fewer than 3 data points (300 ms) were considered to be inconclusive and were merged with the segment before and after them. As previously described (Verdeny-Vilanova et al., 2017), very short segments (with a total displacement <350 nm) that were categorized as active were not considered for run length or velocity analysis. This cut-off displacement of 350 nm was chosen as endolysosmes move at an average speed of ~1-1.7 μm/s and, in 3 frames (300 ms), the expected displacement is ~300-510 nm. The run length and average speed were obtained from the remaining trajectory segments categorized as active.

## Acknowledgements

We thank Prof. Neale D. Ridgway, Dalhousie University for the Hela cells that are a knockout for ORP1L and for the ORP1L WT-mCherry and ORP1L SBD-mCherry constructs. We thank Prof. Michael S. Marks, UPenn, for critical reading and feedback on the manuscript. ML acknowledges funding from the National Institutes of Health/National Institutes for General Medical Sciences (NIH/NIGMS) under the grant numbers: RO1 GM 133842-01 and 1RM1GM136511-01 and the Center for Engineering and Mechanobiology (CEMB) an NSF Science and Technology Center Pilot Award under grant agreement CMMI: 15-48571.

## Author Contributions

ML and ST conceived of the study. ST prepared samples, carried out experiments, wrote software and analyzed data. P.K.R. wrote software and implemented the renormalized voronoi area distribution and KL-Divergence analysis methodology. E.M.S. maintained cell lines, carried out western blot experiments, carried out all transfections, provided reagents and helped with sample preparation. M.T.G. helped carry out dilution and calibration experiments. ML wrote the manuscript, acquired funding and supervised the work. All authors provided feedback on the manuscript.

## References

Abdul-Hammed, M., Breiden, B., Adebayo, M.A., Babalola, J.O., Schwarzmann, G., and Sandhoff, K. (2010). Role of endosomal membrane lipids and NPC2 in cholesterol transfer and membrane fusion. J Lipid Res 51, 1747–1760.

Andronov, L., Orlov, I., Lutz, Y., Vonesch, J.L., and Klaholz, B.P. (2016). ClusterViSu, a method for clustering of protein complexes by Voronoi tessellation in super-resolution microscopy. Sci Rep 6, 24084.

Arumugam, S., and Kaur, A. (2017). The Lipids of the Early Endosomes: Making Multimodality Work. Chembiochem 18, 1053–1060.

Balint, S., Verdeny Vilanova, I., Sandoval Alvarez, A., and Lakadamyali, M. (2013). Correlative live-cell and superresolution microscopy reveals cargo transport dynamics at microtubule intersections. Proc Natl Acad Sci U S A 110, 3375–3380.

Bates, M., Huang, B., Dempsey, G.T., and Zhuang, X. (2007). Multicolor super-resolution imaging with photo-switchable fluorescent probes. Science 317, 1749–1753.

Beatty, W.L. (2006). Trafficking from CD63-positive late endocytic multivesicular bodies is essential for intracellular development of Chlamydia trachomatis. J Cell Sci 119, 350–359.

Belyy, V., Schlager, M.A., Foster, H., Reimer, A.E., Carter, A.P., and Yildiz, A. (2016). The mammalian dynein-dynactin complex is a strong opponent to kinesin in a tug-of-war competition. Nat Cell Biol 18, 1018–1024.

Bissig, C., and Gruenberg, J. (2013). Lipid sorting and multivesicular endosome biogenesis. Cold Spring Harb Perspect Biol 5, a016816.

Brown, C.L., Maier, K.C., Stauber, T., Ginkel, L.M., Wordeman, L., Vernos, I., and Schroer, T.A. (2005). Kinesin-2 is a motor for late endosomes and lysosomes. Traffic 6, 1114–1124.

Brugger, B., Sandhoff, R., Wegehingel, S., Gorgas, K., Malsam, J., Helms, J.B., Lehmann, W.D., Nickel, W., and Wieland, F.T. (2000). Evidence for segregation of sphingomyelin and cholesterol during formation of COPI-coated vesicles. J Cell Biol 151, 507–518.

Cabukusta, B., and Neefjes, J. (2018). Mechanisms of lysosomal positioning and movement. Traffic 19, 761–769.

Cardoso, C.M., Groth-Pedersen, L., Hoyer-Hansen, M., Kirkegaard, T., Corcelle, E., Andersen, J.S., Jaattela, M., and Nylandsted, J. (2009). Depletion of kinesin 5B affects lysosomal distribution and stability and induces peri-nuclear accumulation of autophagosomes in cancer cells. PLoS One 4, e4424.

Cella Zanacchi, F., Manzo, C., Magrassi, R., Derr, N.D., and Lakadamyali, M. (2019). Quantifying Protein Copy Number in Super Resolution Using an Imaging-Invariant Calibration. Biophys J 116, 2195–2203.

Chowdary, P.D., Kaplan, L., Che, D.L., and Cui, B. (2018). Dynamic Clustering of Dyneins on Axonal Endosomes: Evidence from High-Speed Darkfield Imaging. Biophys J 115, 230–241.

Chowdhury, S., Ketcham, S.A., Schroer, T.A., and Lander, G.C. (2015). Structural organization of the dynein-dynactin complex bound to microtubules. Nat Struct Mol Biol 22, 345–347.

Ehmann, N., van de Linde, S., Alon, A., Ljaschenko, D., Keung, X.Z., Holm, T., Rings, A., DiAntonio, A., Hallermann, S., Ashery, U., et al. (2014). Quantitative super-resolution imaging of Bruchpilot distinguishes active zone states. Nature Communications 5, 4650.

Elshenawy, M.M., Canty, J.T., Oster, L., Ferro, L.S., Zhou, Z., Blanchard, S.C., and Yildiz, A. (2019). Cargo adaptors regulate stepping and force generation of mammalian dynein-dynactin. Nat Chem Biol 15, 1093–1101.

Elshenawy, M.M., Kusakci, E., Volz, S., Baumbach, J., Bullock, S.L., and Yildiz, A. (2020). Lis1 activates dynein motility by modulating its pairing with dynactin. Nat Cell Biol 22, 570–578.

Encalada, S.E., Szpankowski, L., Xia, C.H., and Goldstein, L.S. (2011). Stable kinesin and dynein assemblies drive the axonal transport of mammalian prion protein vesicles. Cell 144, 551–565.

Ferro, L.S., Can, S., Turner, M.A., ElShenawy, M.M., and Yildiz, A. (2019). Kinesin and dynein use distinct mechanisms to bypass obstacles. Elife 8.

Franke, C., Repnik, U., Segeletz, S., Brouilly, N., Kalaidzidis, Y., Verbavatz, J.M., and Zerial, M. (2019). Correlative single-molecule localization microscopy and electron tomography reveals endosome nanoscale domains. Traffic 20, 601–617.

Friedman, J.R., Dibenedetto, J.R., West, M., Rowland, A.A., and Voeltz, G.K. (2013). Endoplasmic reticulum-endosome contact increases as endosomes traffic and mature. Mol Biol Cell 24, 1030–1040.

Fu, M.M., and Holzbaur, E.L. (2014). Integrated regulation of motor-driven organelle transport by scaffolding proteins. Trends Cell Biol 24, 564–574.

Fu, M.M., Nirschl, J.J., and Holzbaur, E.L.F. (2014). LC3 binding to the scaffolding protein JIP1 regulates processive dynein-driven transport of autophagosomes. Dev Cell 29, 577–590.

Gould, G.W., and Lippincott-Schwartz, J. (2009). New roles for endosomes: from vesicular carriers to multi-purpose platforms. Nat Rev Mol Cell Biol 10, 287–292.

Granger, E., McNee, G., Allan, V., and Woodman, P. (2014). The role of the cytoskeleton and molecular motors in endosomal dynamics. Semin Cell Dev Biol 31, 20–29.

Guardia, C.M., Farias, G.G., Jia, R., Pu, J., and Bonifacino, J.S. (2016). BORC Functions Upstream of Kinesins 1 and 3 to Coordinate Regional Movement of Lysosomes along Different Microtubule Tracks. Cell Rep 17, 1950–1961.

Hendricks, A.G., Perlson, E., Ross, J.L., Schroeder, H.W., 3rd, Tokito, M., and Holzbaur, E.L. (2010). Motor coordination via a tug-of-war mechanism drives bidirectional vesicle transport. Curr Biol 20, 697–702.

Hu, Y.B., Dammer, E.B., Ren, R.J., and Wang, G. (2015). The endosomal-lysosomal system: from acidification and cargo sorting to neurodegeneration. Transl Neurodegener 4, 18.

Huotari, J., and Helenius, A. (2011). Endosome maturation. EMBO J 30, 3481–3500.

Hyttinen, J.M., Niittykoski, M., Salminen, A., and Kaarniranta, K. (2013). Maturation of autophagosomes and endosomes: a key role for Rab7. Biochim Biophys Acta 1833, 503–510.

Johansson, M., Lehto, M., Tanhuanpaa, K., Cover, T.L., and Olkkonen, V.M. (2005). The oxysterol-binding protein homologue ORP1L interacts with Rab7 and alters functional properties of late endocytic compartments. Mol Biol Cell 16, 5480–5492.

Johansson, M., Rocha, N., Zwart, W., Jordens, I., Janssen, L., Kuijl, C., Olkkonen, V.M., and Neefjes, J. (2007). Activation of endosomal dynein motors by stepwise assembly of Rab7-RILP-p150Glued, ORP1L, and the receptor betalll spectrin. J Cell Biol 176, 459–471.

Keyomarsi, K. (1996). Synchronization of mammalian cells by Lovastatin. Methods in Cell Science 18, 109–114.

Kimura, S., Noda, T., and Yoshimori, T. (2008). Dynein-dependent movement of autophagosomes mediates efficient encounters with lysosomes. Cell Struct Funct 33, 109–122.

Klumperman, J., and Raposo, G. (2014). The complex ultrastructure of the endolysosomal system. Cold Spring Harb Perspect Biol 6, a016857.

Kobayashi, T., Beuchat, M.H., Chevallier, J., Makino, A., Mayran, N., Escola, J.M., Lebrand, C., Cosson, P., Kobayashi, T., and Gruenberg, J. (2002). Separation and characterization of late endosomal membrane domains. J Biol Chem 277, 32157–32164.

Korolchuk, V.I., and Rubinsztein, D.C. (2011). Regulation of autophagy by lysosomal positioning. Autophagy 7, 927–928.

Korolchuk, V.I., Saiki, S., Lichtenberg, M., Siddiqi, F.H., Roberts, E.A., Imarisio, S., Jahreiss, L., Sarkar, S., Futter, M., Menzies, F.M., et al. (2011). Lysosomal positioning coordinates cellular nutrient responses. Nat Cell Biol 13, 453–460.

Lemmon, M.A. (2008). Membrane recognition by phospholipid-binding domains. Nat Rev Mol Cell Biol 9, 99–111.

Levet, F., Hosy, E., Kechkar, A., Butler, C., Beghin, A., Choquet, D., and Sibarita, J.B. (2015). SR-Tesseler: a method to segment and quantify localization-based super-resolution microscopy data. Nat Methods 12, 1065–1071.

Maday, S., Twelvetrees, A.E., Moughamian, A.J., and Holzbaur, E.L. (2014). Axonal transport: cargo-specific mechanisms of motility and regulation. Neuron 84, 292–309.

McKenney, R.J., Huynh, W., Tanenbaum, M.E., Bhabha, G., and Vale, R.D. (2014). Activation of cytoplasmic dynein motility by dynactin-cargo adapter complexes. Science 345, 337–341.

Mohan, N., Sorokina, E.M., Verdeny, I.V., Alvarez, A.S., and Lakadamyali, M. (2019). Detyrosinated microtubules spatially constrain lysosomes facilitating lysosome-autophagosome fusion. J Cell Biol 218, 632–643.

Nirschl, J.J., Magiera, M.M., Lazarus, J.E., Janke, C., and Holzbaur, E.L. (2016). alpha-Tubulin Tyrosination and CLIP-170 Phosphorylation Regulate the Initiation of Dynein-Driven Transport in Neurons. Cell Rep 14, 2637–2652.

Olenick, M.A., and Holzbaur, E.L.F. (2019). Dynein activators and adaptors at a glance. J Cell Sci 132.

Olkkonen, V.M., and Li, S. (2013). Oxysterol-binding proteins: sterol and phosphoinositide sensors coordinating transport, signaling and metabolism. Prog Lipid Res 52, 529–538.

Perez-Cruz, F. (2008). Kullback-Leibler divergence estimation of continuous distributions. Paper presented at: IEEE international symposium on information theory.

Pfeffer, S.R. (2001). Rab GTPases: specifying and deciphering organelle identity and function. Trends Cell Biol 11, 487–491.

Pu, J., Guardia, C.M., Keren-Kaplan, T., and Bonifacino, J.S. (2016). Mechanisms and functions of lysosome positioning. J Cell Sci 129, 4329–4339.

Puchner, E.M., Walter, J.M., Kasper, R., Huang, B., and Lim, W.A. (2013). Counting molecules in single organelles with superresolution microscopy allows tracking of the endosome maturation trajectory. Proc Natl Acad Sci U S A 110, 16015–16020.

Rai, A., Pathak, D., Thakur, S., Singh, S., Dubey, A.K., and Mallik, R. (2016). Dynein Clusters into Lipid Microdomains on Phagosomes to Drive Rapid Transport toward Lysosomes. Cell 164, 722–734.

Reck-Peterson, S.L., Redwine, W.B., Vale, R.D., and Carter, A.P. (2018). The cytoplasmic dynein transport machinery and its many cargoes. Nat Rev Mol Cell Biol 19, 382–398.

Redpath, G.M.I., Betzler, V.M., Rossatti, P., and Rossy, J. (2020). Membrane Heterogeneity Controls Cellular Endocytic Trafficking. Front Cell Dev Biol 8, 757.

Rocha, N., Kuijl, C., van der Kant, R., Janssen, L., Houben, D., Janssen, H., Zwart, W., and Neefjes, J. (2009). Cholesterol sensor ORP1L contacts the ER protein VAP to control Rab7-RILP-p150 Glued and late endosome positioning. J Cell Biol 185, 1209–1225.

Rosa-Ferreira, C., and Munro, S. (2011). Arl8 and SKIP act together to link lysosomes to kinesin-1. Dev Cell 21, 1171–1178.

Schroeder, C.M., and Vale, R.D. (2016). Assembly and activation of dynein-dynactin by the cargo adaptor protein Hook3. J Cell Biol 214, 309–318.

Soppina, V., Rai, A.K., Ramaiya, A.J., Barak, P., and Mallik, R. (2009). Tug-of-war between dissimilar teams of microtubule motors regulates transport and fission of endosomes. Proc Natl Acad Sci U S A 106, 19381–19386.

Tanemura, M. (2003). Statistical distributions of Poisson Voronoi cells in two and three dimensions. FORMA-TOKYO 18, 221–247.

Torres, S., Balboa, E., Zanlungo, S., Enrich, C., Garcia-Ruiz, C., and Fernandez-Checa, J.C. (2017). Lysosomal and Mitochondrial Liaisons in Niemann-Pick Disease. Front Physiol 8, 982.

Urnavicius, L., Lau, C.K., Elshenawy, M.M., Morales-Rios, E., Motz, C., Yildiz, A., and Carter, A.P. (2018). Cryo-EM shows how dynactin recruits two dyneins for faster movement. Nature 554, 202–206.

van der Kant, R., Zondervan, I., Janssen, L., and Neefjes, J. (2013). Cholesterol-binding molecules MLN64 and ORP1L mark distinct late endosomes with transporters ABCA3 and NPC1. J Lipid Res 54, 2153–2165.

Vanlandingham, P.A., and Ceresa, B.P. (2009). Rab7 regulates late endocytic trafficking downstream of multivesicular body biogenesis and cargo sequestration. J Biol Chem 284, 12110–12124.

Verdeny-Vilanova, I., Wehnekamp, F., Mohan, N., Sandoval Alvarez, A., Borbely, J.S., Otterstrom, J.J., Lamb, D.C., and Lakadamyali, M. (2017). 3D motion of vesicles along microtubules helps them to circumvent obstacles in cells. J Cell Sci 130, 1904–1916.

Vihervaara, T., Uronen, R.L., Wohlfahrt, G., Bjorkhem, I., Ikonen, E., and Olkkonen, V.M. (2011). Sterol binding by OSBP-related protein 1L regulates late endosome motility and function. Cell Mol Life Sci 68, 537–551.

Wijdeven, R.H., Janssen, H., Nahidiazar, L., Janssen, L., Jalink, K., Berlin, I., and Neefjes, J. (2016). Cholesterol and ORP1L-mediated ER contact sites control autophagosome transport and fusion with the endocytic pathway. Nat Commun 7, 11808.

Zanacchi, F.C., Manzo, C., Alvarez, A.S., Derr, N.D., Garcia-Parajo, M.F., and Lakadamyali, M. (2017). A DNA origami platform for quantifying protein copy number in super-resolution. Nat Methods 14, 789–792.

Zhao, K., and Ridgway, N.D. (2017). Oxysterol-Binding Protein-Related Protein 1L Regulates Cholesterol Egress from the Endo-Lysosomal System. Cell Rep 19, 1807–1818.

