## Supplemental Figures for "ORP1L regulates dynein clustering on endolysosmal membranes in response to cholesterol levels"

### Supplementary Figures:

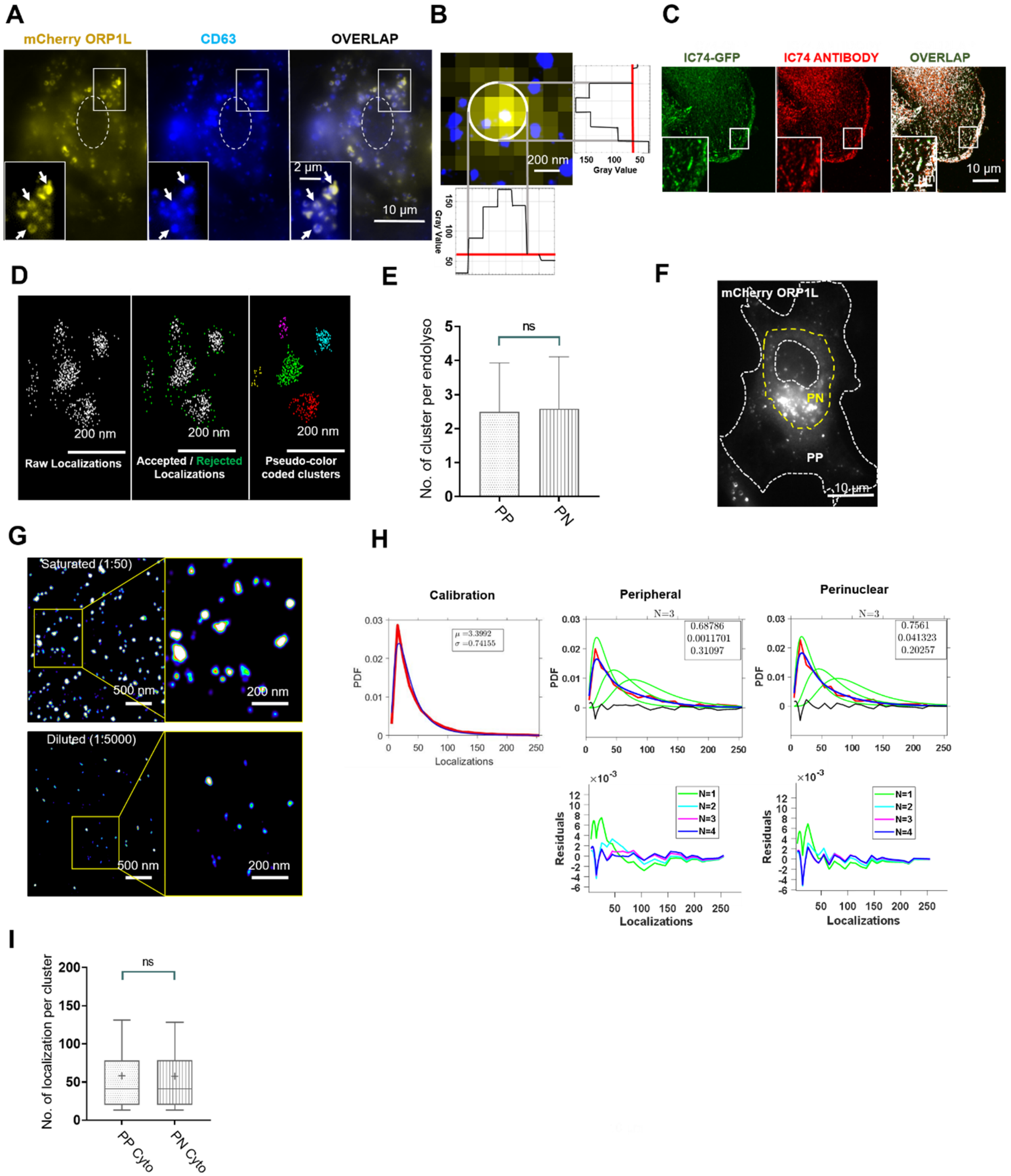


**Figure S1:** Analysis of dynein copy number on endolysosomal compartments.

(A) Widefield image of full length ORP1L (ORP1L WT-mCherry, *left*, yellow), immunofluorescence image of CD63 (*middle*, blue) and overlay (*right*) in ORP1L mCherry-

WT expressing Hela ORP1L-KO cells. The nucleus is shown with white dashed lines. A zoom of the region inside the white square is shown as inset. Arrows point to some examples of mCherry-ORP1L-WT and CD63 double positive endolysosomes.

(B) A representative conventional image of an individual endolysosome (yellow) overlaid with the super-resolution image of dynein (blue) in ORP1L WT-mCherry expressing Hela ORP1L-KO cells. The plot shows the intensity profile of the endolysosome along the x- and y-axis. To identify dynein clusters overlapping with ORP1L positive compartments, the intensity profile of conventional ORP1L image was used as a mask. The line intensity profile of the ORP1L image along the x- and y-axis was plotted using image J, and the width across one third of the full intensity maxima was considered as the mask (gray lines and white circle).

(C) Wide field image of dynein IC74-GFP (*left*, green), immunofluorescence image of dynein IC74 (*middle*, red) and overlay (*right*, white) in Hela cells stably expressing dynein IC74 fused to GFP. A zoom of the region inside the white square is shown as inset.

(D) (*Left*) Raw super-resolution localization data for dynein. Each white dot corresponds to the position of a localized fluorophore. Fluorophore positions (localizations) are used as seeds for Voronoi tessellation such that a Voronoi polygon is drawn around each localization. The size of the Voronoi polygons corresponds to localization density such that clustered localizations having a higher density will also have smaller Voronoi polygons. (*Middle*) A threshold is imposed such that Voronoi polygons having a size smaller than the threshold are accepted (white) and those having a size larger than the threshold are rejected (green). (*Right*) The accepted localizations are segmented into individual nano-clusters. Nano-clusters are pseudo color-coded.

(E) Bar plot showing the average number of dynein nano-clusters per endolysosome in peripheral (PP) (*n=96 endolysosomes from n=6 cells, n=2experiments*, mean = 2.5±1.4) and perinuclear (PN) (*n=64 endolysosomes from n=6 cells, n=2 experiments*, mean = 2.6±1.5) regions of ORP1L WT-mCherry expressing Hela ORP1L-KO cells, nano-clusters were obtained after segmentation of dynein super-resolution images using Voronoi tessellation. The whiskers correspond to standard deviation. Statistical significance was assessed using a Kolmogorov-Smirnov -test with a p-value of > 0.999.

(F) Wide-field image of ORP1L WT-mCherry in Hela ORP1L-KO cells. The cell edge and the nucleus are shown with white dashed lines. The region corresponding to the cell peri-nuclear area is shown with yellow dashed lines.

(G) Super-resolution image of dynein nano-clusters in Hela WT cells labeled with 1:5000 dilution of the primary antibody to sparsely label single IC74 subunits (*bottom*) or 1:50 dilution of the primary antibody to saturate labeling of dynein motors (*top*). The secondary antibody dilution was the same under the two conditions.

(H) Calibration (*Left top*): Plot showing the number of localizations per dynein nano-cluster under dilute labeling condition (red) and the log normal fit (blue) to obtain the calibration function ($f_1$) for single IC74 with fit parameters $\mu = 3.39$ and $\sigma = 0.74$. (*Middle and right top*): Plot showing the number of localizations per dynein nano-cluster for peripheral/peri-nuclear endolysosomes under normal (experimental) labeling conditions (red); dimeric ($f_2$), trimeric ($f_3$) calibration functions obtained from a linear convolution of the monomeric calibration function (i.e. the log normal fit, $f_1$) (green); and the fit of the experimental data to a combination of $f_1$, $f_2$ and $f_3$ (blue) with weights corresponding to 1, 2 and 3 IC74s as indicated in the box inset. (*Middle and right bottom*) Residuals calculated after fitting $f_1$ (green, N=1), combination of $f_1$ and $f_2$ (blue, N=2), combination of $f_1$, $f_2$, $f_3$ (magenta, N=3) and combination of $f_1$, $f_2$, $f_3$, $f_4$ (indigo, N=4) to peripheral/peri-nuclear experimental data.

(I) Box plot showing the number of localizations per cytosolic dynein nano-clusters from peripheral regions (PP Cyto) (from n=6 cells, n=2experiments, mean 58±50), and perinuclear regions (PN Cyto) (from n=6 cells, n=2 experiments, mean 57.5±49.5) of ORP1L WT-mCherry expressing Hela ORP1L-KO cells. The box corresponds to 25-75 percentile, the line corresponds to the median, the cross corresponds to the mean, and the whiskers correspond to 10-90 percentile. Statistical significance was assessed using a Kolmogorov-Smirnov-test with a p-value of 0.7.

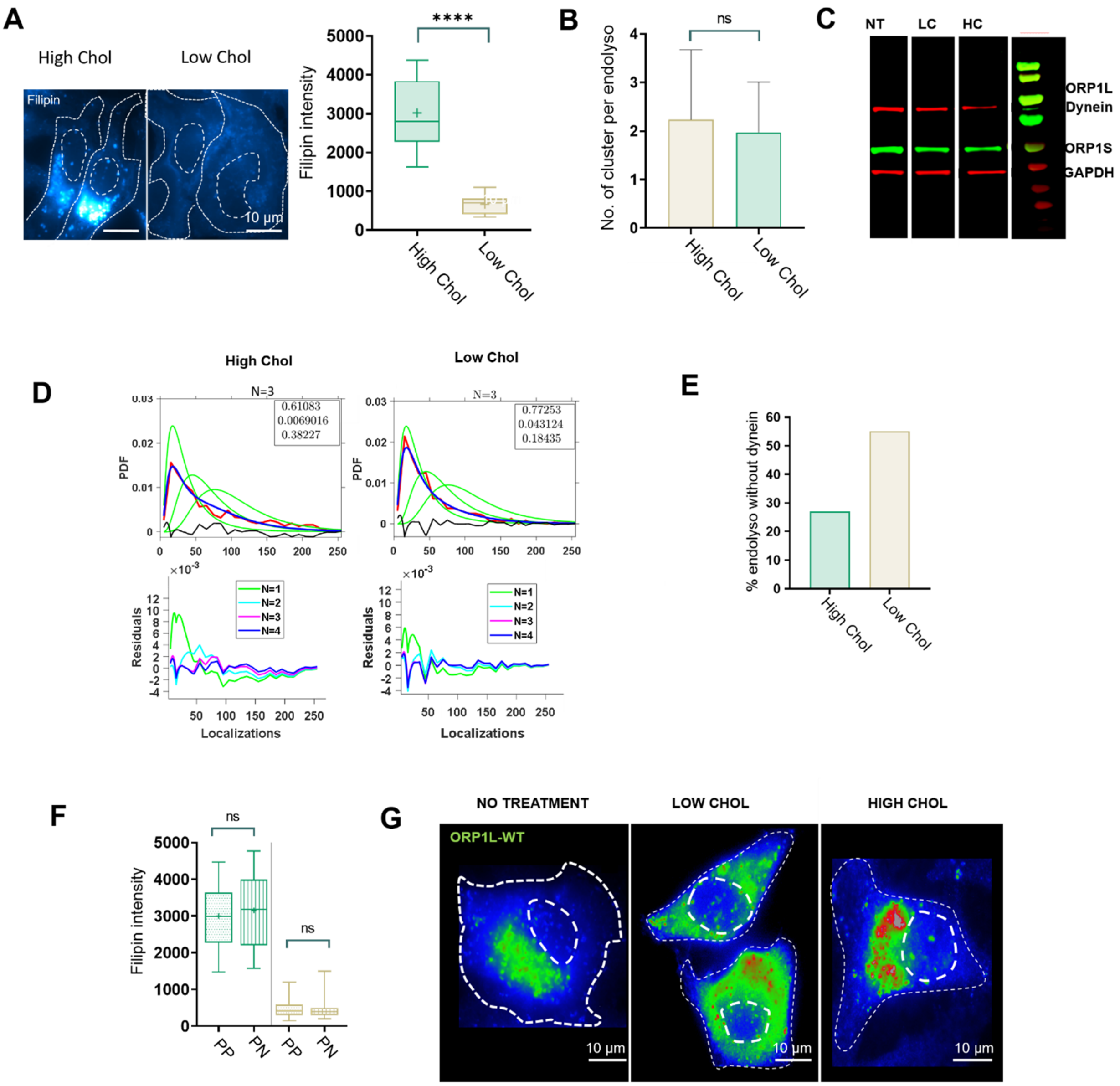


**Figure S2:** Cholesterol levels regulate endolysosomal positioning

(A) *(Left)* Wide-field images showing Filipin labelled endolysosmes in ORP1L WT-mCherry expressing Hela ORP1L-KO cells treated with U18666A (High Chol) and Lovastatin (Low Chol). A color lookup table shows the intensities with the same contrast settings. *(Right)* Box plot showing the filipin intensity of endolysosomal compartments in these cells treated with U18666A (High Chol) (*n=224 endolysosomes from n=6 cells, n=2 experiments, mean 71±1.9)*) and Lovastatin (Low Chol) (*n=217 endolysosomes from n=6 cells, n=2 experiments, mean* 50±1.9). The box corresponds to 25-75 percentile, the line corresponds to the median, the cross corresponds to the mean and the whiskers correspond to 10-90 percentile. Statistical significance was assessed using a Kolmogorov-Smirnov-test with a p-value of <0.0001.

(B) Bar plot showing the average number of dynein nano-clusters per endolysosome in ORP1L WT-mCherry expressing Hela ORP1L-KO cells treated with U18666A (High Chol) (*n=70 endolysosomes from n=6 cells, n=2 experiments*, mean = 2±1) and Lovastatin (Low Chol)l (*n=77 endolysosomes from n=6 cells, n=2experiments*, mean = 2.2±1.4), clusters were obtained after segmentation of dynein super-resolution images using Voronoi tessellation. The whiskers correspond to standard deviation. Statistical significance was assessed using a Kolmogorov-Smirnov -test with a p-value of 0.6.

(C) Western blot showing dynein intermediate chain (red) and the long (ORP1L) and short (ORP1S) isoforms of ORP1 (green) in HeLa ORPIL-KO cells. Housekeeping gene GAPDH is shown in red. First lane corresponds to untreated cells (NT), second lane to cells treated for 6 hours with Lovastatin (LC) and third lane to cells treated for 12 hours with U18666A (HC). The lanes were cropped from a larger gel containing additional, not relevant conditions.

(D) (*Top*) Plot showing the number of localizations per dynein nano-cluster for endolysosomes in cells treated with U18666A (High Chol) or Lovastatin (Low Chol) under normal (experimental) labeling conditions (red); dimeric ($f_2$), trimeric ($f_3$) calibration functions obtained from a linear convolution of the monomeric calibration function (i.e. the log normal fit, $f_1$) (green); and the fit of the experimental data to a combination of $f_1$, $f_2$ and $f_3$ (blue) with weights corresponding to 1, 2, and 3 IC74s as indicated in the box inset. (*Bottom*) Residuals calculated after fitting $f_1$ (green, N=1), combination of $f_1$ and $f_2$ (blue, N=2), combination of $f_1$, $f_2$, $f_3$ (magenta, N=3) and combination of $f_1$, $f_2$, $f_3$, $f_4$ (indigo, N=4) to High and Low Cholesterol experimental data.

(E) Bar plot showing the percentage of endolysosomes without any dynein nano-cluster in ORP1L WT-mCherry expressing Hela ORP1L-KO cells treated with U18666A (High Chol: 27%) (n=83 endolysosomes from n=6 cells, n=2 experiments) and Lovastatin (Low Chol: 55%) (n=265 endolysosomes from n=6 cells, n=2 experiments).

(F) Box plot showing the filipin intensity of peripherally (PP) positioned and peri-nculearly (PN) positioned endolysosomal compartments in ORP1L WT-mCherry expressing Hela

ORP1L-KO cells treated with U18666A (High Chol) (n=50 endolysosomes from n=4 cells, n=2 experiments, mean = 2998±209 a.u. for PP and n=30 endolysosomes from n=4 cells, n=2 experiments, mean = 3152±180 a.u. for PN) and Lovastatin (Low Chol) (n=40 endolysosomes from n=4 cells, n=2 experiments mean = 491±55 a.u. for PP and n=50 endolysosomes from n=4 cells, n=2 experiments, mean 480±43 a.u. for PN). The box corresponds to 25-75 percentile, the line corresponds to the median, the cross corresponds to the mean and the whiskers correspond to 10-90 percentile. Statistical significance was assessed using a Kolmogorov-Smirnov-test with a p-value of 0.91 and 0.99.

(G) Wide-field images showing ORP1L WT-mCherry labelled endolysosomes (expressed in Hela ORPL1-KO cells) in untreated cells (*left*), cells treated with Lovastatin (*middle*, Low Chol) and cells treated with U18666A (*right*, High Chol).

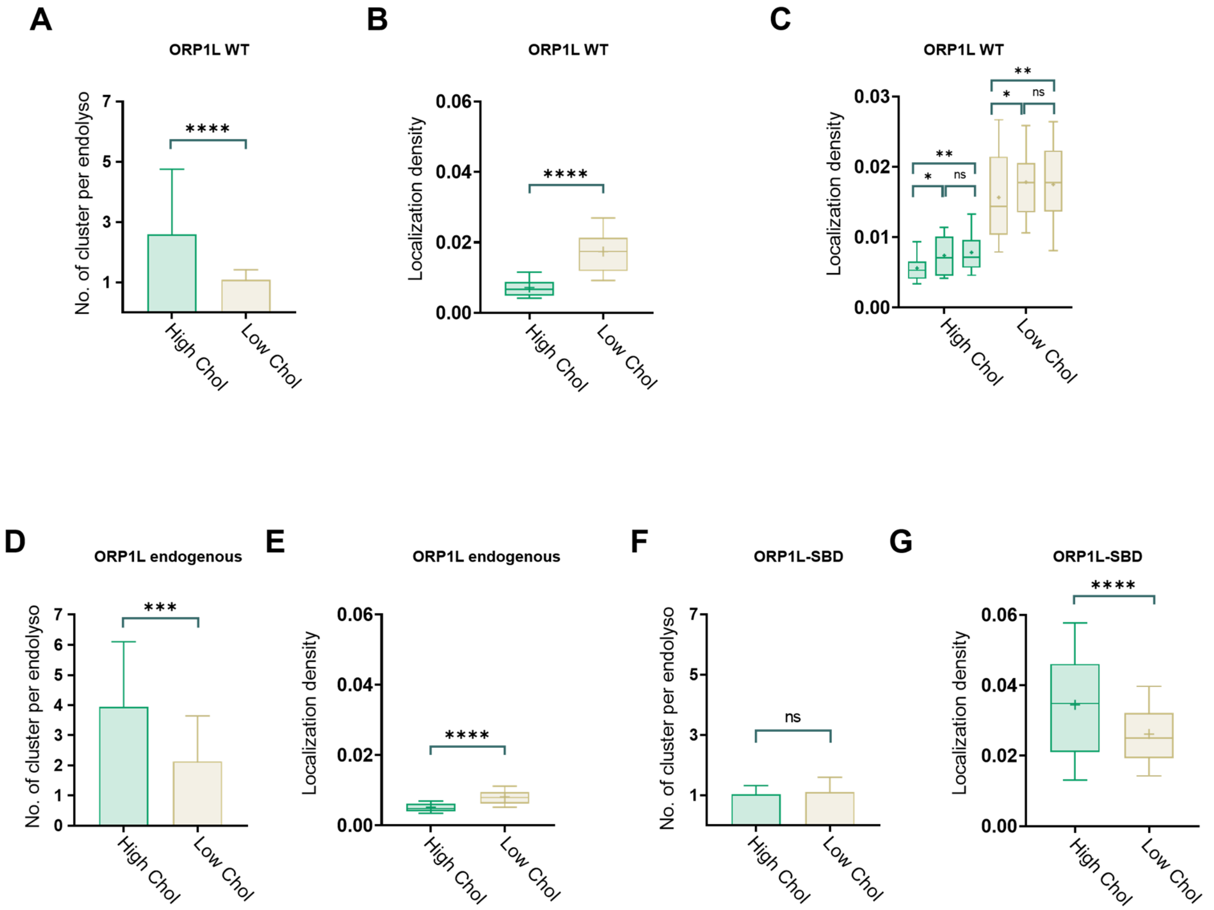


**Figure S3:** ORP1L is more clustered on endolysosomes with high cholesterol content

(A) Bar plot showing the average number of ORP1L clusters per endolysosome in U18666A (High Chol) (n=110 endolysosomes from n=6 cells, n=2 experiments, mean = 2.6±0.32) and Lovastatin (Low Chol) (n=120 endolysosomes from n=6 cells, n=2 experiments, mean = 1±0.04) treated ORP1L WT-mCherry expressing Hela ORP1L-KO cells after segmentation of ORP1L super-resolution images using Voronoi tessellation. The whiskers correspond to standard deviation. Statistical significance was assessed using a Kolmogorov-Smirnov-test with a p-value of <0.0001.

(B) Box plot showing the total number of localizations per endolysosomal unit area calculated from ORP1L super-resolution images in ORP1L WT-mCherry expressing Hela ORP1L-KO cells treated with U18666A (High Chol) (n=110 endolysosomes from n=6 cells, n=2 experiments, mean = 0.0072±0.00032) and Lovastatin (Low Chol) (n=120 endolysosomes from n=6 cells, n=2 experiments, mean = 0.017±0.00062). The box corresponds to 25-75 percentile, the line corresponds to the median, the cross corresponds to the mean and the whiskers correspond to 10-90 percentile. Statistical significance was assessed using a Kolmogorov-Smirnov-test with a p-value of <0.0001.

(C) (High Chol) Box plots showing the total number of localizations per endolysosomal unit area calculated from three independent biological replicates of ORP1L super-resolution images of endolysosomes in ORP1L WT-mCherry expressing Hela ORP1L-KO cells treated with U18666A (High Chol) (Box 1: n=22 endolysosomes from n=5 cells, n=1 experiment, mean = 0.006±0.002; Box 2: n=27 endolysosomes from n=2 cells, n=1 experiments, mean = 0.007±0.003; Box 3: n=32 endolysosomes from n=2 cells, n=1 experiments, mean = 0.008±0.003). (Low Chol) Same as High Chol, but in cells treated with Lovastatin (Box 1: n=29 endolysosomes from n=5 cells, n=1 experiment, mean = 0.016±0.007; Box 2: n=42 endolysosomes from n=2 cells, n=1 experiments, mean = 0.018±0.006; Box 3: n=36 endolysosomes from n=2 cells, n=1 experiments, mean = 0.017±0.003). Statistical significance was assessed using a Kolmogorov-Smirnov-test with a p-value of (High Chol) *: 0.049, ns: 0.62, **: 0.01 and (Low Chol) ns: 0.33, ns: 0.82, ns: 0.47.

(D) Bar plot showing the average number of endogenous ORP1L clusters per endolysosome in HeLa cells treated with U18666A (High Chol) (n=60 endolysosomes from n=5 cells, n=2 experiments, mean = 4±0.31) and Lovastatin (Low Chol) (n=65 endolysosomes from n=5 cells, n=2 experiments, mean = 2±0.02) after segmentation of ORP1L super-resolution images using Voronoi tessellation. The whiskers correspond to standard deviation.

Statistical significance was assessed using a Kolmogorov-Smirnov-test with a p-value of 0.0002.

(E) Box plot showing the total number of localizations per endolysosomal unit area calculated from endogenous ORP1L super-resolution images in HeLa cells treated with U18666A (High Chol) (n=60 endolysosomes from n=5 cells, n=2 experiments, mean = 0.0052±0.0002) and Lovastatin (Low Chol) (n=65 endolysosomes from n=5 cells, n=2 experiments, mean = 0.008±0.0003). The box corresponds to 25-75 percentile, the line corresponds to the median, the cross corresponds to the mean and the whiskers correspond to 10-90 percentile. Statistical significance was assessed using a Kolmogorov-Smirnov-test with a p-value of <0.0001.

(F) Bar plot showing the average number of ORP1L clusters per endolysosome in ORP1L SBD-mCherry expressing Hela ORP1L-KO cells treated with U18666A (High Chol) (n=120 endolysosomes from n=6 cells, n=2 experiments, mean = 1±0.03 and 1.1±0.04) and Lovastatin (Low Chol) (n=170 endolysosomes from n=6 cells, n=2 experiments, mean = 1.1±0.04) after segmentation of ORP1L super-resolution images using Voronoi tessellation. The whiskers correspond to standard deviation. Statistical significance was assessed using a Kolmogorov-Smirnov -test with a p-value of 0.992.

(G) Box plot showing the total number of localizations per endolysosomal unit area calculated from ORP1L super-resolution images in ORP1L SBD-mCherry expressing cells treated with U18666A (High Chol) (n=120 endolysosomes from n=6 cells, n=2 experiments, mean = 0.034±0.0017) and Lovastatin (Low Chol) (n=170 endolysosomes from n=6 cells, n=2 experiments, mean = 0.026±0.00075). The box corresponds to 25-75 percentile, the line corresponds to the median, the cross corresponds to the mean and the whiskers correspond to 10-90 percentile. Statistical significance was assessed using a Kolmogorov-Smirnov-test with a p-value of <0.0001.

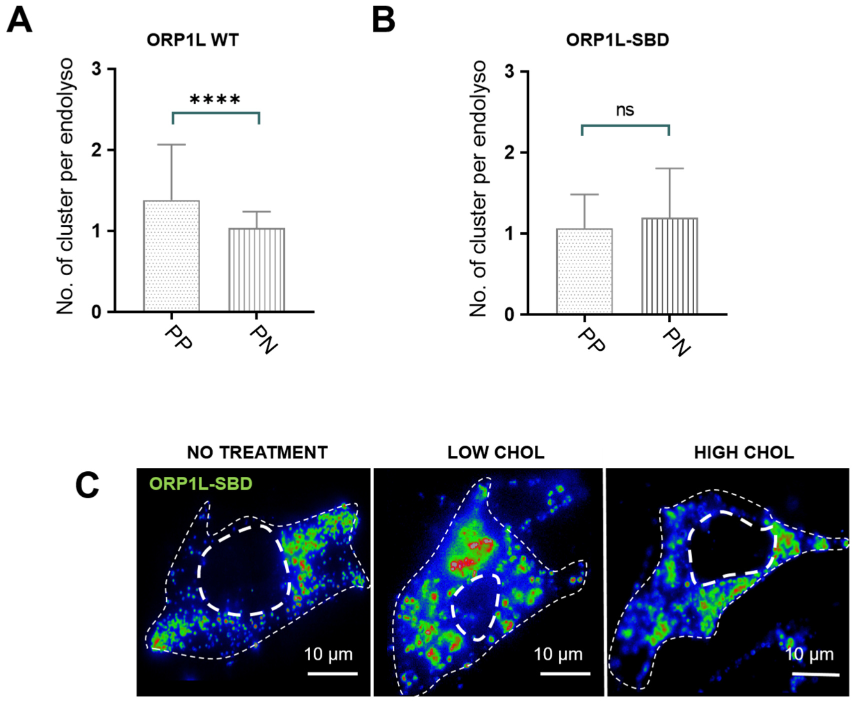


**Figure S4:** ORP1L clustering and endolysosome positioning are dependent on ORP1L's cholesterol binding domain

(A) Bar plot showing the average number of ORP1L clusters per endolysosome for peripherally positioned (PP) endolysosomes (n=140 endolysosomes from n=6 cells, n=2 experiments, mean = 1.4±0.05) and perinuclearly positioned endolysosomes (PN) (n=200 endolysosomes from n=6 cells, n=2 experiments, mean = 1±0.01) in untreated ORP1L WT-mCherry expressing Hela ORP1L-KO cells after segmentation of ORP1L super-resolution images using Voronoi tessellation. The whiskers correspond to standard deviation. Statistical significance was assessed using a Kolmogorov-Smirnov-test with a p-value of <0.0001.

(B) Bar plot showing the number of ORP1L clusters per endolysosome for peripherally positioned (PP) endolysosomes (n=150 endolysosomes from n=6 cells, n=2 experiments, mean = 1.1±0.033) and perinuclearly positioned endolysosomes (PN) (n=160 endolysosomes from n=6 cells, n=2 experiments, mean = mean = 1.1±0.05) in untreated cells expressing ORP1L SBD-mCherry after segmentation of ORP1L super-resolution images using Voronoi tessellation. The whiskers correspond to standard deviation. Statistical significance was assessed using a Kolmogorov-Smirnov -test with a p-value of 0.47.

(C) Wide-field images of ORP1L SBD-mCherry labelled endolysosomal compartments in untreated cells (left), cells treated with Lovastatin (middle, Low Chol) and cells treated with U18666A (right, High Chol).
